# Precision-based causal inference modulates audiovisual temporal recalibration

**DOI:** 10.1101/2024.03.08.584189

**Authors:** Luhe Li, Fangfang Hong, Stephanie Badde, Michael S. Landy

## Abstract

Cross-modal temporal recalibration guarantees stable temporal perception across ever-changing environments. Yet, the mechanisms of cross-modal temporal recalibration remain unknown. Here, we conducted an experiment to measure how participants’ temporal perception was affected by exposure to audiovisual stimuli with consistent temporal delays. Consistent with previous findings, recalibration effects plateaued with increasing audiovisual asynchrony and varied by which modality led during the exposure phase. We compared six observer models that differed in how they update the audiovisual temporal bias during the exposure phase and whether they assume modality-specific or modality-independent precision of arrival latency. The causal-inference observer shifts the audiovisual temporal bias to compensate for perceived asynchrony, which is inferred by considering two causal scenarios: when the audiovisual stimuli have a common cause or separate causes. The asynchrony-contingent observer updates the bias to achieve simultaneity of auditory and visual measurements, modulating the update rate by the likelihood of the audiovisual stimuli originating from a simultaneous event. In the asynchrony-correction model, the observer first assesses whether the sensory measurement is asynchronous; if so, she adjusts the bias proportionally to the magnitude of the measured asynchrony. Each model was paired with either modality-specific or modality-independent precision of arrival latency. A Bayesian model comparison revealed that both the causal-inference process and modality-specific precision in arrival latency are required to capture the nonlinearity and asymmetry observed in audiovisual temporal recalibration. Our findings support the hypothesis that audiovisual temporal recalibration relies on the same causal-inference processes that govern cross-modal perception.

## 1 Introduction

Perception is not rigid but rather can adapt to the environment. In a multimodal environment, misalignment across the senses can occur because signals in different modalities may arrive with different physical and neural delays in the relevant brain areas (Fain, 2019; Pöppel, 1988; Spence & Squire, 2003). Perceptual misalignment can also arise from changes in the perceptual system relative to the environment, such as when wearing a virtual reality headset or adapting to hearing aids. Cross-modal temporal recalibration serves as a critical mechanism to maintain perceptual synchrony despite changes in the perceptual systems and the environment (reviewed in King, 2005; Vroomen and Keetels, 2010). This phenomenon is exemplified in audiovisual temporal recalibration, where consistent exposure to audiovisual stimulus-onset asynchrony (SOA) shifts the point of subjective simultaneity between auditory and visual stimuli; as a result, stimuli perceived as temporally discrepant at first are gradually perceived as more synchronous (Di Luca et al., 2009; Fujisaki et al., 2004; Hanson et al., 2008; Harrar & Harris, 2008; Heron et al., 2007; Keetels & Vroomen, 2007; Navarra et al., 2005; Roach et al., 2011; Tanaka et al., 2011; Vatakis et al., 2007, 2008; Vroomen & de Gelder, 2004; Vroomen & Keetels, 2010).

However, the mechanisms of cross-modal temporal recalibration remain unknown. The current models of audiovisual temporal recalibration either did not specify the recalibration process (Di Luca et al., 2009; Navarra et al., 2009; Yarrow et al., 2015), or cannot fully capture the characteristics of recalibration effects (Roach et al., 2011; Sato & Aihara, 2011; Yarrow et al., 2015). Specifically, audiovisual temporal recalibration shows two distinct characteristics: the amount of recalibration is nonlinear and asymmetric as a function of the SOA participants are adapted to (adapter SOA). The amount of recalibration is not proportional to the adapter SOA, but instead plateaus at an SOA of approximately 100–300 ms (Fujisaki et al., 2004; Vroomen & de Gelder, 2004). Recalibration can also be asymmetrical: the magnitude of recalibration differs when the visual stimulus leads during the exposure phase compared to when the auditory stimulus leads (Fujisaki et al., 2004; O’Donohue et al., 2022; Van der Burg et al., 2013). These observations can provide insights into the mechanisms of cross-modal temporal recalibration.

Here, we propose a causal-inference model to explain the mechanism of audiovisual temporal recalibration. Causal inference is the process in which the observer determines whether multisensory signals originate from a common source and should be integrated or kept separate (Sato et al., 2007; Shams & Beierholm, 2010; Wei & Körding, 2009). Bayesian models based on causal inference have been proposed to explain multisensory integration effects (Körding et al., 2007; Sato et al., 2007), and these models have been empirically validated in studies of spatial audiovisual and visual-tactile integration (Badde, Navarro, & Landy, 2020; Beierholm et al., 2009; Rohe & Noppeney, 2015; Wozny et al., 2010). In the temporal domain, some studies have successfully used causal inference to model the integration of cross-modal relative timing, accurately predicting simultaneity judgments in audiovisual speech (Magnotti et al., 2013) and more complex scenarios involving one auditory and two visual stimuli (Sato, 2021).

In the context of cross-modal recalibration, causal inference is expected to play a role based on the intuition that recalibration should be reduced when the multisensory signals are not perceived as causally related (Fujisaki et al., 2004; Hsiao et al., 2022; Vroomen & de Gelder, 2004). Supporting this, causal-inference models successfully predicted cross-modal spatial recalibration of visual-auditory (Hong, 2023; Hong et al., 2021; Sato et al., 2007) and visuo-tactile (Badde, Navarro, & Landy, 2020) signals. Building on this framework, here we propose a causal-inference model for cross-modal temporal recalibration that derives the multisensory percept based on inferences about the shared origin of the signals and updates the cross-modal temporal biases such that subsequent measurements are shifted toward the percept.

The first aim of this study is to test whether performing causal inference is necessary to explain the nonlinearity of audiovisual temporal recalibration across different adapter SOAs. To this aim, we compared the causal-inference model with two alternatives: an asynchrony-contingent model and an asynchrony-correction model. The asynchrony-contingent model scales the amount of recalibration by the likelihood that the sensory measurement of SOA was caused by a synchronous audiovisual stimulus pair. The model predicts a nonlinear recalibration effect across adapter SOAs without requiring observers to perform full Bayesian inference. The asynchrony-correction model assumes that recalibration only occurs when an asynchronous onset of the cross-modal stimuli is registered, followed by the update of the cross-modal temporal bias to compensate for this SOA measurement. This account is based on the intuitive rationale that repeated measurements of asynchrony can prompt the perceptual system to restore coherence. In contrast, this model predicts minimal recalibration when the adapter SOA falls within the range of measured asynchronies that can arise with simultaneously presented stimuli due to sensory noise. This model serves as the baseline for model comparison.

The second aim was to examine factors that had the potential to drive the asymmetry of recalibration across visual-leading and auditory-leading adapter SOAs. It has been suggested that the asymmetry may be explained by physical and neural latency differences between signals (O’Donohue et al., 2022; Van der Burg et al., 2013). These latency differences can vary significantly based on the physical distance between the stimulus and the sensors, as well as the neural transmission time required for the signal to reach the relevant sensory region (Badde, Navarro, & Landy, 2020; Hirsh & Sherrick, 1961; King, 2005). While these latency differences can explain the audiovisual temporal bias observed in most humans, they would affect recalibration to different adapter SOAs equally, making it unlikely for any asymmetry to arise. In contrast to latency differences, sensory uncertainty has been shown to affect the degree of cross-modal recalibration in a complex fashion (Badde, Navarro, & Landy, 2020; Hong et al., 2021; van Beers et al., 2002). We hypothesized that the difference across modalities in the variability of the arrival times, the time it takes visual and auditory signals to arrive in the relevant brain areas, plays a critical role in the asymmetry of cross-modal temporal recalibration.

To examine the mechanism underlying audiovisual temporal recalibration, we manipulated the adapter SOA cross sessions, introducing asynchronies up to 0.7 s of either auditory or visual lead. Before and after the exposure phase in each session, we measured participants’ perception of audiovisual relative timing using a ternary temporal-order-judgement (TOJ) task. To preview the empirical results, we confirmed the nonlinearity of the recalibration effect: recalibration magnitude increased linearly for short adapter SOAs, but then reached an asymptote or even decreased with increasing adapter SOAs. Furthermore, participants showed idiosyncratic asymmetries of the recalibration effect across modalities; for most participants, the amount of recalibration was larger when the auditory stimulus led than when it lagged, but the opposite was found for other participants. To scrutinize the factors that might drive the nonlinearity and asymmetry of temporal recalibration, we fitted six models to the data. These models based the amount of recalibration either on perceptual causal-inference processes, a heuristic evaluation of the common cause of the audiovisual stimuli, or a fixed criterion for the need to correct asynchrony. For each of these three models we implemented either modality-specific or modality-independent precision of the arrival times. The model comparison revealed that the assumptions of Bayesian causal inference combined with modality-specific precision are essential to accurately capture the nonlinearity and idiosyncratic asymmetry of temporal recalibration.

## 2 Results

### 2.1 Behavioral results

We adopted a classical three-phase recalibration paradigm in which participants completed a pre-test, an exposure phase, and a post-test in each session. In pre- and post-tests, we measured participants’ perception of audiovisual relative timing using a ternary TOJ task: participants reported the perceived order (“visual first,” “auditory first,” or “simultaneous”) of audiovisual stimulus pairs with varying SOA (range: from -0.5 to 0.5 s with 15 levels; Figure 1A). In the exposure phase, we induced temporal recalibration by having participants perform a control task, the oddball-detection task. Specifically, participants were exposed to a series of audiovisual stimuli with a consistent SOA (250 trials; Figure 1B). To ensure that participants were attentive to the stimuli, we inserted oddball stimuli with greater intensity in either one or both modalities (5% of the total trials independently sampled for each modality). Participants were instructed to press a key corresponding to the auditory, visual, or both oddballs whenever an oddball stimulus appeared. The high *d*^*′*^ of oddball-detection performance (auditory *d*^*′*^ = 3.34*±*0.54, visual *d*^*′*^ = 2.44*±*0.72) indicates that participants paid attention to both modalities. The post-test was almost identical to the pre-test, except that before every temporal-order-judgment trial, there were three top-up oddball-detection trials to maintain the recalibration effect. In total, participants completed nine sessions on separate days. The adapter SOA (range: -0.7 to 0.7 s) was fixed within a session, but varied randomly across sessions and participants.

**Figure 1:**
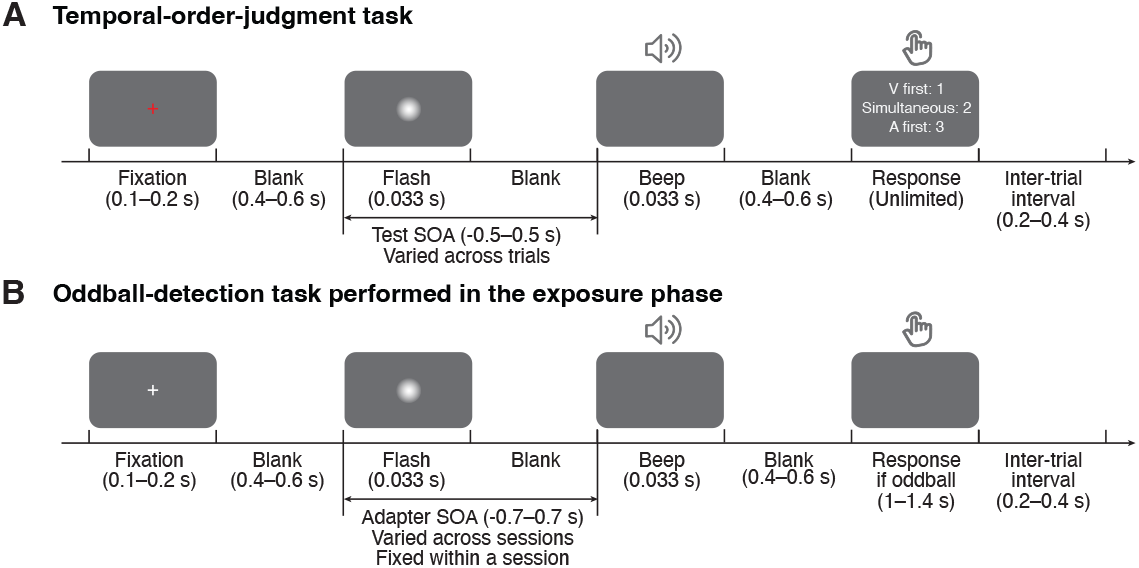
Task timing. (A) Temporal-order-judgment task administered in the pre- and post-tests. In each trial, participants made a temporal-order judgment in response to an audiovisual stimulus pair with a varying stimulus-onset asynchrony (SOA). Negative values: auditory lead; positive values: visual lead. The contrast of the visual stimulus has been increased for this illustration. (B) Oddball-detection task performed in the exposure phase and top-up trials during the post-exposure test phase. Participants were repeatedly presented with an audiovisual stimulus pair with a SOA that was fixed within each session but varied across sessions. Occasionally, the intensity of either one or both of the stimuli was increased. Participants were instructed to press a key corresponding to the auditory, visual, or both oddballs whenever an oddball stimulus appeared.

We compared the temporal-order judgments between the pre- and post-tests to examine the amount of audiovisual temporal recalibration induced by the audiovisual stimuli during the exposure phase. Specifically, we fitted the data from the pre- and post-tests jointly assuming different points of subjective simultaneity (PSS) between the two tests while assuming the same shape for the psychometric functions that is determined by the relative arrival-latencies, their precision, and fixed response criteria (Figure 2A; see Supplement Section 1 for the formalization of the atheoretical model and an alternative model assuming a shift in the response criteria due to recalibration). The PSS is the physical SOA that corresponds to the maximum probability of reporting simultaneity (Sternberg & Knoll, 1973). The amount of audiovisual temporal recalibration was defined as the difference between the two PSS’s. At the group level, we observed a nonlinear pattern of recalibration as a function of the adapter SOA: the amount of recalibration in the direction of the adapter SOA first increased but then plateaued with increasing magnitude of the adapter SOA, the SOA of the pairs presented during the exposure phase (Figure 2B). Additionally, we observed an asymmetry in the amount of recalibration between auditory-leading and visual-leading adapter SOAs, with auditory-leading adapter SOAs inducing a greater amount of recalibration (Figure 2B; see Supplement Figure S2A for individual participants’ data). To quantify this asymmetry for each participant, we calculated an asymmetry index, defined as the sum of the recalibration effects across all adapter SOAs (zero: no evidence for asymmetry; positive values: greater recalibration given visual-lead adapters; negative: greater recalibration given auditory-lead adapters). For each participant, we bootstrapped the temporal-order judgments to obtain a 95% confidence interval for the asymmetry index. Eight out of nine participants showed an asymmetry index significantly different from zero, with the majority showing greater recalibration for auditory-leading adapter SOAs, suggesting a general asymmetry in recalibration (Supplement Figure S2B).

**Figure 2:**
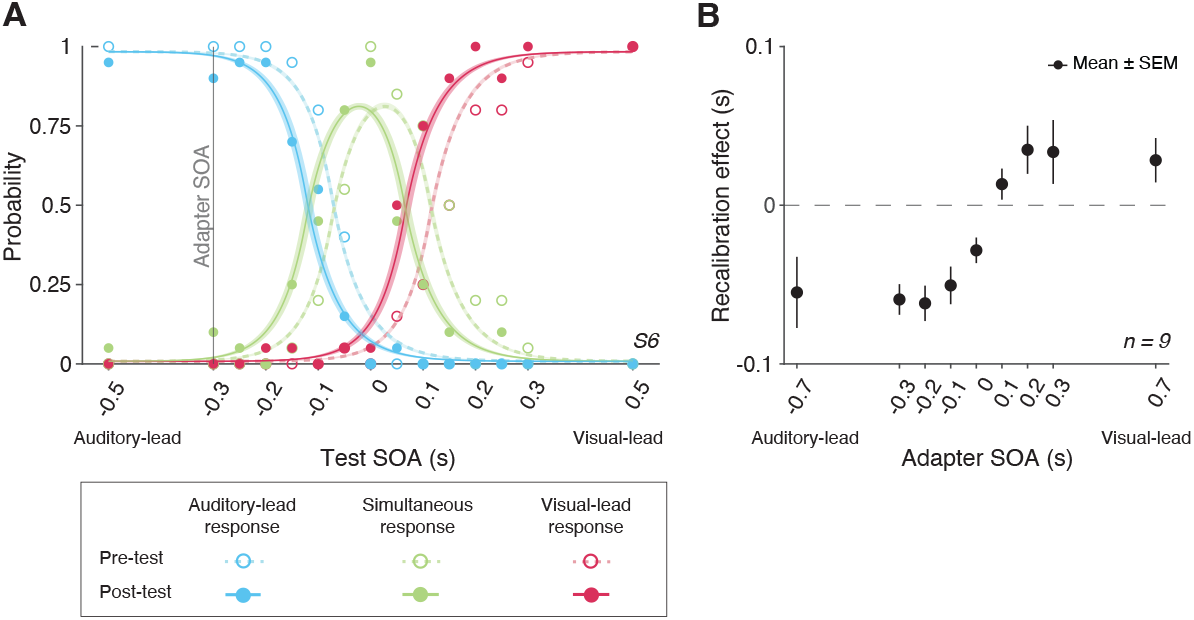
Behavioral results. (A) The probability of reporting that the auditory stimulus came first (blue), the two arrived at the same time (green), or the visual stimulus came first (red) as a function of SOA for a representative participant in a single session. The adapter SOA was -0.3 s for this session. Curves: best-fitting psychometric functions estimated jointly using the data from the pre-test (dashed) and post-test (solid). Shaded areas: 95% bootstrapped confidence intervals. (B) Mean recalibration effects averaged across all participants as a function of adapter SOA. The recalibration effects are defined as the shifts in the point of subjective simultaneity (PSS) from the pre-to the post-test, where the PSS is the physical SOA at which the probability of reporting simultaneity is maximized. Error bars: *±*SEM.

### 2.2 Modeling results

In the following sections, we describe our models for cross-modal temporal recalibration by first laying out the general assumptions of these models, and then elaborating on the differences between them. Then, we compare the models’ ability to capture the observed data.

#### 2.2.1 General model assumptions

We formulated six process models of cross-modal temporal recalibration (Figure 3). These models share several assumptions about audiovisual temporal perception and recalibration that we selected based on a comparison of atheoretical, descriptive models of our data (Supplement Section 1). First, when an auditory and a visual signal are presented, the corresponding neural signals arrive in the relevant brain areas with a variable latency due to internal and external noise. We assume arrival times for the two modalities are independent and that the arrival latencies are exponentially distributed (García-Pérez & Alcalá-Quintana, 2012) (Figure 3A, left panel). Moreover, we assume a constant offset between auditory and visual arrival times, reflecting an audiovisual temporal bias. A simple derivation shows that the resulting measurement of SOA has a double-exponential distribution (Figure 3A, right panel; see derivation in Supplement Section 3). The probability density function peaks at a SOA that is the physical SOA of the stimuli plus the participant’s audiovisual temporal bias. The slopes of the measurement distribution reflect the precision of the arrival times; the steeper the slope, the more precise the measured latency. When the precision differs between modalities, the measurement distribution of the SOA between the auditory and visual stimuli is asymmetrical (Figure 3B).

**Figure 3:**
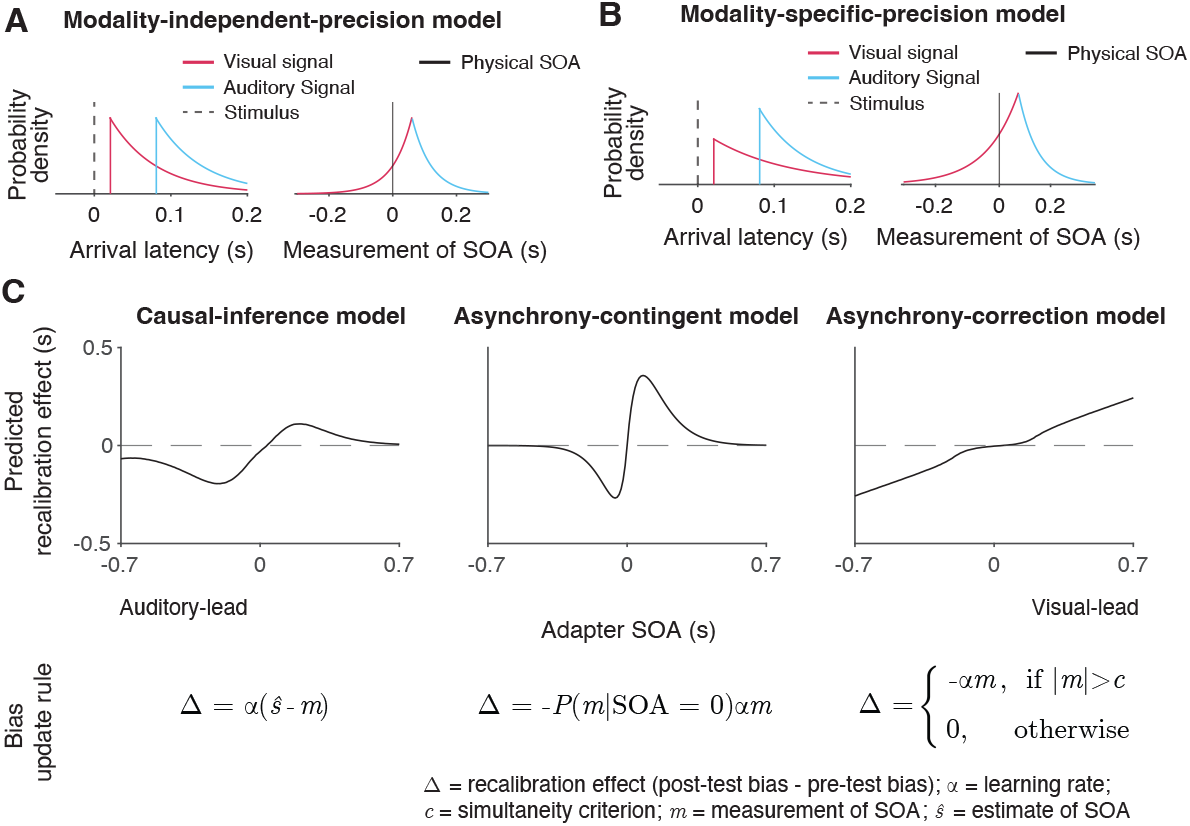
Illustration of the six observer models of cross-modal temporal recalibration. (A) Left: Arrival-latency distributions for auditory (blue) and visual (red) sensory signals. When the precision of arrival latency is modality-independent, these two exponential distributions have identical shape. Right: The resulting symmetrical double-exponential measurement distribution of the SOA of the stimuli. (B) When the precision of the arrival latencies is modality-dependent, the arrival-latency distributions for auditory and visual signals have different shapes, and the resulting measurement distribution of the SOA is asymmetrical. (C) Bias update rules and predicted recalibration effects for the three contrasted recalibration models: The causal-inference model updates the audiovisual bias based on the difference between the estimated and measured SOA. The asynchrony-contingent model updates the audiovisual bias by a proportion of the measured SOA and modulates the update rate by the likelihood that the measured sensory signals originated from a simultaneous audiovisual pair. The asynchrony-correction model adjusts the audiovisual bias by a proportion of the measured SOA when this measurement exceeds fixed critera for simultaneity.

Second, these models define temporal recalibration as the accumulation of updates to the audiovisual temporal bias after each encounter with an SOA. The accumulated shift in the audiovisual bias at the end of the exposure phase is then carried over to the post-test phase and persists throughout. Lastly, the bias is assumed to be reset to the same initial value in the pre-test across all nine sessions, reflecting the stability of the audiovisual temporal bias over time (Grabot & van Wassenhove, 2017).

#### 2.2.2 Models of cross-modal temporal recalibration

The six models we tested differed in the mechanism governing the updates of the audiovisual bias during the exposure phase as well as the modality-specificity of the precision of arrival times.

We formulated a temporal variant of the spatial Bayesian causal-inference model of recalibration (Badde, Navarro, & Landy, 2020; Hong, 2023; Hong et al., 2021; Sato et al., 2007) to describe the recalibration of the relative timing between cross-modal stimuli (Figure 3C, left panel). In this model, when an observer is presented with an audiovisual stimulus pair during the exposure phase, they compute two intermediate estimates of the SOA between the stimuli, one for the common-cause scenario and the other for the separate-cause scenario. In the common-cause scenario, the estimated SOA of the stimuli is smaller than the measured SOA as it is combined with a prior distribution over SOA that reflects simultaneity. In the separate-causes scenario, the estimated SOA is approximately equal to the measured SOA. The two estimates are then averaged with each one weighted by the posterior probability of the corresponding causal scenario. The audiovisual bias is then updated to reduce the difference between the measured SOA and the combined estimate of the SOA. In other words, causal inference regulates the recalibration process by shifting the measured SOA to more closely match the percept, which in turn is computed based on the inferred causal structure. The asynchrony-contingent model assumes that the observer estimates the likelihood that the sensory signals originated from a simultaneous audiovisual pair and updates the audiovisual bias by a proportion of measured SOA scaled by this likelihood (Figure 3C, middle panel). There is a key distinction between the likelihood of simultaneity and the likelihood of a common cause. The likelihood of a common cause considers the prior distribution of SOAs when signals originate from the same source, including nonzero probabilities for SOAs≠ 0. In contrast, the likelihood of simultaneity exclusively considers the case when SOA = 0. Additionally, we assume that asynchrony-contingent observer computes the likelihood of simultaneity based on the knowledge of the double-exponential measurement distribution, instead of assuming a Gaussian measurement distribution as was done previously (Maij et al., 2009). The update rate of the audiovisual bias is proportional to this likelihood. For a stimulus pair with a large SOA, the average likelihood of the stimuli being physically simultaneous decreases, leading to reduced recalibration effects compared to stimulus pairs with smaller SOAs. Thus, this asynchrony-contingent model is capable of replicating the nonlinearity of recalibration across adapter SOAs without requiring the observer to perform full Bayesian inference.

The asynchrony-correction model assumes that the observer first compares the sensory measurement of SOA to their criteria for audiovisual simultaneity to decide whether to recalibrate in a given trial. If the measured SOA falls within the range perceived as simultaneous according to the fixed criteria, the observer might attribute a non-zero measurement of SOA to sensory noise and omit recalibration. On the other hand, if the measured SOA exceeds this range, the observer perceives the stimuli as asynchronous, and shifts the audiovisual bias by a proportion of the measurement of SOA (Figure 3C, right panel). This model serves as a direct contrast to the causal-inference model, as it predicts an opposite pattern: a nonlinear but monotonic increase in temporal recalibration, with minimal recalibration when the measured SOA falls within the simultaneity range and increasing recalibration as the measured SOA moves further outside of this range.

We additionally assumed either modality-specific or modality-independent precision of the arrival times. Each choice suggests a different origin of the variability. Either the variability of the arrival times is limited by neural-latency noise in each sensory channel (Yarrow et al., 2022) and thus is modality-specific or the variability of arrival times results from the variability in a central timing mechanism (Hirsh & Sherrick, 1961) and is thus modality-independent.

#### 2.2.3 Model fitting and model comparison

We fitted six models to each participant’s data. Each model was constrained jointly by the temporal-order judgments from the pre- and post-tests of all nine sessions. To quantify model performance, we calculated model evidence, i.e., the likelihood of each model given the data marginalized over all possible parameters, which revealed that the causal-inference model had the strongest model evidence at the group level and best fit the data of most participants, followed by the asynchrony-contingent model and then the asynchrony-correction model. To quantify the differences between model performance, we performed a Bayesian model comparison by computing the Bayes factor for each model relative to the worst-performing model, the asynchrony-correction model with modality-independent arrival-latency precision (Figure 4A, see Supplement Figure S3 for individual-level model comparison). Within each of these three model categories, the version incorporating modality-specific precision consistently outper-formed the modality-independent version.

**Figure 4:**
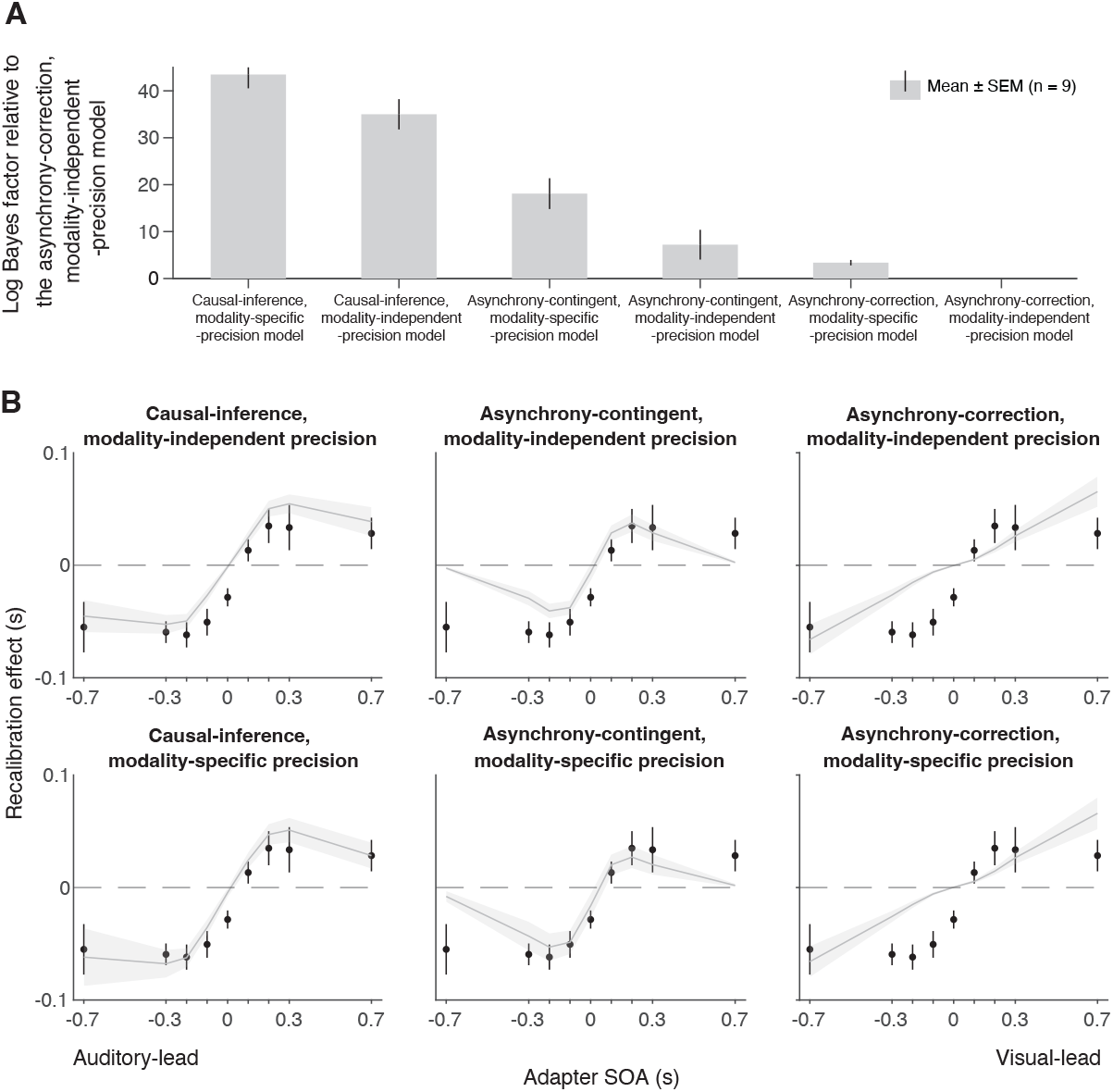
Model comparison and predictions. (A) Model comparison based on model evidence. Each bar represents the group-averaged log Bayes Factor of each model relative to the asynchrony-correction, modality-independent-precision model, which had the weakest model evidence. (B) Empirical data (points) and model predictions (lines and shaded regions) for the recalibration effect as a function of adapter SOA.

#### 2.2.4 Model prediction

To inspect the quality of the model fit, for every model, we used the best-fitting parameter estimates for each participant to predict the group-average recalibration effect as a function of adapter SOA (Figure 4B). The nonlinearity of audiovisual temporal recalibration across adapter SOAs was captured by both the asynchrony-contingent and causal-inference models. Nonetheless, the causal-inference model outperformed the asynchrony-contingent model by accurately predicting a non-zero average recalibration effect at adapter SOAs of 0.7 s and -0.7 s, where the asynchrony-contingent model predicted no recalibration. Additionally, incorporating modality-specific precision enabled both the asynchrony-contingent and causalinference models to more accurately predict increased recalibration when the adapter SOA was auditory-leading. Overall, the model that relies on causal inference during the exposure phase and assumes modality-specific precision of arrival times most accurately captured both the nonlinearity and asymmetry of the recalibration effect. This model could also account for individual participants’ idiosyncratic asymmetry in temporal recalibration to auditory- and visual-leading adapter SOAs (see Supplement Figure S4 for predictions of individual participants’ recalibration effects of all models; see Figure S6 for predictions of individual participants’ TOJ responses using the causal-inference models with modality-specific precision).

#### 2.2.5 Model simulation

Simulations with the causal-inference model revealed which factors of the modeled recalibration process determine the degree of nonlinearity and asymmetry of cross-modal temporal recalibration to different adapter SOAs. The prior belief that the auditory and visual stimuli share a common cause plays a crucial role in adjudicating the relative influence of the two causal scenarios (Figure 5A). When the observer has a prior belief that audiovisual stimuli always originate from the same source, they recalibrate by a proportion of the perceived SOA no matter how large the measured SOA is, mirroring the behavior of the asynchrony-correction model when its criteria for simultaneity are such that no stimuli are treated as simultaneous. On the other hand, when the observer believes that the audiovisual stimuli always have separate causes, they treat the audiovisual stimuli as independent of each other and do not recalibrate. Estimates of the common-cause prior for our participants fall between the two extreme beliefs, resulting in the nonlinear pattern of recalibration that lies between the extremes of no recalibration and the proportional recalibration effects as a function of the adapter SOA (see Supplement Section 6.1 for parameter estimates for individual participants).

**Figure 5:**
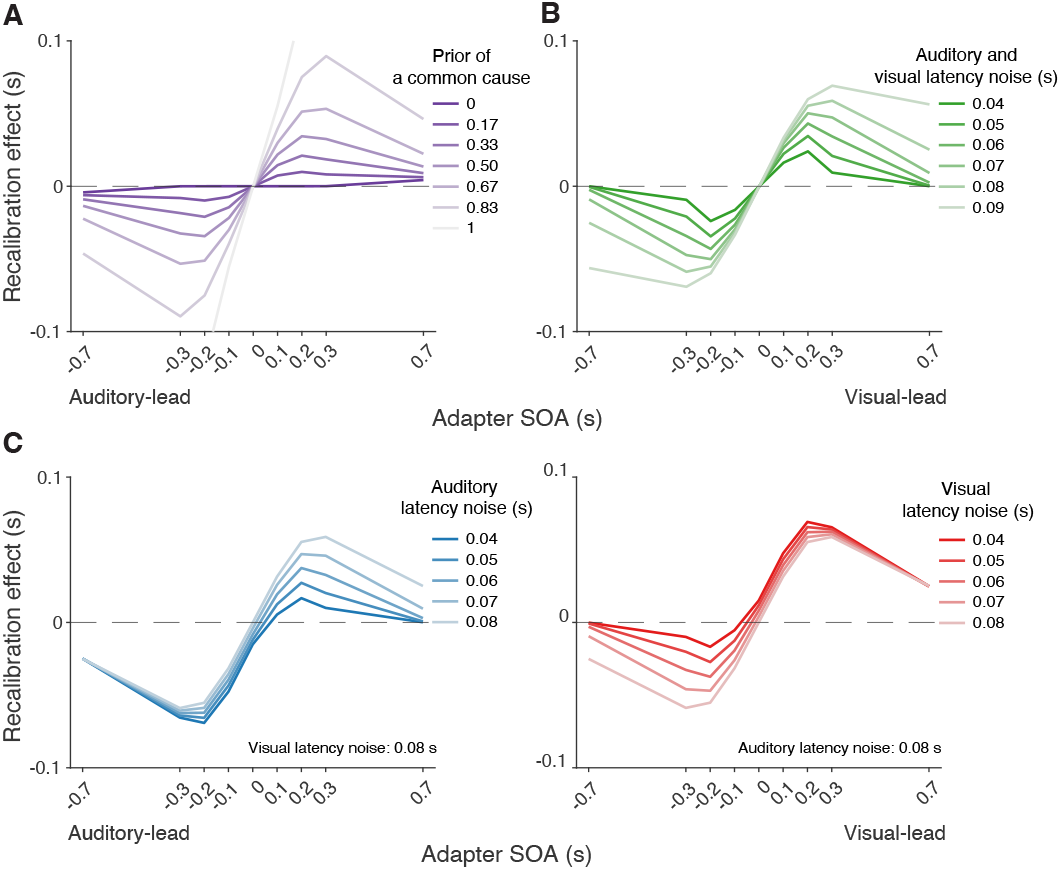
Simulation of temporal recalibration using the causal-inference model. (A) The influence of the observer’s prior assumption of a common cause: the stronger the prior, the larger the recalibration effects. (B) The influence of latency noise: recalibration effects increase with decreasing sensory precision (i.e., increasing latency noise captured by the exponential time constant) of both modalities. (C) The influence of auditory/visual latency noise: recalibration effects are asymmetric between auditory-leading and visual-leading adapter SOAs due to differences in the precision of auditory and visual arrival latencies. Left panel: Increasing auditory latency precision (i.e., reducing auditory latency noise) reduces recalibration in response to visual-leading adapter SOAs. Right panel: Increasing visual precision (i.e., reducing visual latency noise) reduces recalibration in response to auditory-leading adapter SOAs.

Simulations also identified key model elements of the causal-inference model that predict a non-zero recalibration effect even at large SOAs, a feature that distinguishes the causal-inference from the asynchrony-contingent model. This non-zero recalibration effect for large adapter SOAs can be replicated by either assuming a strong prior for a common cause (Figure 5A) or by assuming low sensory precision of arrival times (Figure 5B). Both relationships are intuitive: observers with a stronger prior belief in a common cause and ideal observers with lower sensory precision are more likely to assign a higher posterior probability to the common-cause scenario, leading to greater recalibration. A decrease of the spread of the prior distribution over SOA conditioned on a common cause increases the recalibration magnitude, but only over a small range of SOAs for which there is a higher probability of the common-cause scenario (Supplement Figure S8A), and thus cannot account for non-zero recalibration for large SOAs.

Differences in arrival-time precision between audition and vision result in an asymmetry of audiovisual temporal recalibration across adapter SOAs (Figure 5C). The amount of recalibration is attenuated when the modality with the higher precision lags the less precise one during the exposure phase. When the more recent stimulus component in a cross-modal pair is more precise, the perceptual system is more likely to attribute the asynchrony to separate causes and thus recalibrate less. In addition, the fixed audiovisual bias does not affect asymmetry, but shifts the recalibration function laterally and determines the adapter SOA for which no recalibration occurs (Supplement Figure S8B).

## 3 Discussion

This study scrutinized the mechanism underlying audiovisual temporal recalibration. We measured the effects of exposure to audiovisual stimulus pairs with a constant temporal offset (adapter SOA) on audiovisual temporal-order perception across a wide range of adaptor SOAs. Recalibration effects changed nonlinearly with the magnitude of adapter SOAs and were asymmetric across auditory-leading and visual-leading adapter SOAs. We then compared the predictions of different observer models for the amount of recalibration as a function of adapter SOA. A Bayesian causal-inference model with modality-specific precision of the arrival latencies fit the observed data best. These findings suggest that human observers rely on causal-inference-based percepts to recalibrate cross-modal temporal perception. These results align closely with studies that have demonstrated the role of causal inference in audiovisual (Hong et al., 2021) and visual-tactile spatial recalibration (Badde, Navarro, & Landy, 2020). Our results are also consistent with previous recalibration models that assumed a strong relation between perception and recalibration (Sato, 2021; Sato et al., 2007). Hence, we suggest that the same mechanisms underly cross-modal perception and recalibration across different sensory features.

The observed recalibration results could not be predicted by the asynchrony-contingent model that employed a heuristic approximation of the causal-inference process. Even though this model was capable of predicting a nonlinear relationship between the recalibration effect and the adapter SOA, it failed to capture a non-zero recalibration effect at large adapter SOAs shown by several of our participants. The reason for that is that this model uses the likelihood of a synchronous audiovisual stimulus pair given the measured SOA to modulate the update rate of audiovisual bias, which will be very small on average for large SOAs. Therefore, the model predicts little to no recalibration at large adaptor SOAs. In contrast, the causal-inference model can capture the non-zero recalibration effect because the common-cause scenario always influences the amount of recalibration even when the adapter SOA is too large to be perceived as synchronous. Simulation (Figure 5A, B) shows that a strong prior belief in a common cause or less precision of arrival times can result in non-zero recalibration effects following exposure to clearly asynchronous stimulus pairs. Notably, even though it might at first seem counter-intuitive that cross-modal temporal recalibration can be elicited by clearly asynchronous streams of sensory information, many of us have experienced this effect during laggy, long video conferences.

The asynchrony-correction model assumes that observers recalibrate to restore temporal synchrony whenever the SOA measurement indicates a temporal discrepancy, but this model predicts recalibration effects across adapter SOAs that are contrary to our observations. This suggests that cross-modal temporal recalibration is not merely triggered by an asynchronous sensory measurement of SOA and an attempt to correct it. In contrast, the causal-inference model accurately captured the plateau of the recalibration effects as adapter SOA increased, because the probability that the auditory and visual stimuli have separate causes also increased. This resulted in a smaller discrepancy between the sensory measurement and the final percept of the SOA, leading to less recalibration.

We found that most of our participants exhibited larger recalibration effects in response to exposure to audiovisual stimuli with a consistent auditory lead compared to exposure to a visual lead. This result is consistent with a previous study that reported greater cumulative recalibration in response to audiovisual stimuli with an auditory-lead at the group level (O’Donohue et al., 2022). Our simulation results further suggested that this asymmetry in recalibration effects might be due to higher precision of auditory compared to visual arrival latencies. A few participants displayed the opposite pattern: stronger recalibration effects following exposure to visual-leading audiovisual stimuli. This is not surprising, as causalinference models often reveal substantial individual differences in sensory noise (Hong et al., 2021; Magnotti et al., 2013). A recent EEG study further provided neural correlates for individual sensory noise by identifying correlations between neural-latency noise and behavioral sensory noise measured from simultaneity-judgment tasks for audiovisual, visuo-tactile, and audio-tactile pairs (Yarrow et al., 2022). Therefore, our model explains how individual differences in precision of arrival latency could contribute to the asymmetry in cross-modal temporal recalibration observed in previous studies. For example, Fujisaki et al. (2004) found a slightly larger recalibration in response to audiovisual stimuli with a visual lead compared to an auditory lead, while their pilot results with the same design but a wider range of adapter SOAs showed the opposite pattern.

In order to incorporate causal inference in our recalibration models, we modeled recalibration as a shift of audiovisual bias. Building on previous latency-shift models (Di Luca et al., 2009; Navarra et al., 2009), we specified a mechanism for how the audiovisual bias is updated during the exposure to an audiovisual SOA. Our model is not mutually exclusive with other models that implement recalibration as a shift of simultaneity criteria (Yarrow, Jahn, et al., 2011; Yarrow et al., 2015), or a change of sensitivity to discriminate SOA (Roseboom et al., 2015). A possible implementation of recalibration at the circuity level is given by models assuming that audiovisual offsets are encoded by populations of neurons tuned to different SOAs. In these models, recalibration is the consequence of selective gain reduction of neurons tuned to SOAs similar to the adapter SOA (Cai et al., 2012; Roach et al., 2011; Yarrow et al., 2015). Simulations show that this model can predict nonlinear recalibration effects as a function of adapter SOA depending on the number of neurons and the range of preferred SOAs (Supplementary Section S8). However, to capture the asymmetric recalibration effects depending on which modality leads, one needs to incorporate inhomogenous neuronal selectivity, i.e., unequal tuning curves, for auditory-leading and visual-leading SOAs.

Causal inference may effectively function as a credit-assignment mechanism to enhance perceptual accuracy during recalibration. In sensorimotor adaptation, humans correct motor errors that are more likely attributed to their own motor system rather than to the environment (Berniker & Kording, 2008; Wei & Körding, 2009). In visuomotor adaptation, substantial temporal recalibration occurs in response to exposure to movement-leading SOAs but less so to visual-leading SOAs (Rohde & Ernst, 2012; Rohde et al., 2014), because only movement-leading SOAs can be interpreted as causally linked sensory feedback from a preceding movement.

Causal-inference-based recalibration can further solve the conundrum that humans, despite our ability for cross-modal temporal recalibration, show persistent temporal biases (Grabot & van Wassenhove, 2017). These audiovisual and visual-tactile temporal biases appear to be shaped by early sensory experience (Badde, Ley, et al., 2020) and seem to be resistant to recalibration. The persistence of these biases contradicts recalibration models that reduce the measured cross-modal asynchrony. Instead, our causal-inference-based models of recalibration include an assumption that recalibration eliminates the discrepancy between measured and inferred asynchrony, both of which are influenced by cross-modal biases.

Previous studies have probed the role of causal inference for temporal recalibration and perception by experimentally varying task-irrelevant cues to a shared origin of the crossmodal stimuli, with mixed results. Earlier studies found no significant change in temporal recalibration when altering the sound presentation method (headphones versus a speaker) or switching the presentation ear (Fujisaki et al., 2004), nor did recalibration effects vary with the spatial alignment of the audiovisual stimulus pair (Keetels & Vroomen, 2007). However, subsequent studies provide evidence that spatial grouping influences temporal recalibration, with the PSS shifting toward the temporal relationship suggested by spatially co-located stimuli (Heron et al., 2012; Yarrow, Roseboom, & Arnold, 2011). Others found that spatial cues (Heron et al., 2012; Yuan et al., 2012) and featural content cues such as male or female audiovisual speech and high-pitch sounds (Roseboom & Arnold, 2011; Roseboom et al., 2013; Yuan et al., 2012) are both determinants of cross-modal temporal recalibration. Recent studies on audiovisual integration have extended causal-inference models to account for both the spatial position and temporal discrepancy of audiovisual signals (Hong, 2023; McGovern et al., 2016). Conversely, perceived conflicts in task-irrelevant features of visual-haptic stimuli do not influence the integration of task-relevant features, suggesting that causal inference is feature-specific rather than pertaining to whole objects (Badde et al., 2023). These studies suggest that task-relevant spatial and temporal information are taken into account for causal inference and might play a critical role in cross-modal recalibration.

## 4 Methods

### 4.1 Participants

Ten students from New York University (three males; age: 24.4 *±* 1.77; all right-handed) participated in the experiment. They all reported normal or corrected-to-normal vision. All participants provided informed written consent before the experiment and received $15/hr as monetary compensation. The study was conducted in accordance with the guidelines laid down in the Declaration of Helsinki and approved by the New York University institutional review board. One out of ten participants was identified as an outlier and therefore excluded from further data analysis (Supplement Figure S9).

### 4.2 Apparatus and stimuli

Participants completed the experiments in a dark and semi sound-attenuated room. They were seated 1 m from an acoustically transparent, white screen (1.36 × 1.02 m, 68 × 52° visual angle) and placed their head on a chin rest. An LCD projector (Hitachi CP-X3010N, 1024 × 768 pixels, 60 Hz) was mounted above and behind participants to project visual stimuli on the screen. The visual and auditory stimulus durations were 33.33 ms. The visual stimulus was a high-contrast (36.1 cd/m^2^) Gaussian blob (SD: 3.6°) on a gray background (10.2 cd/m^2^) projected onto the screen. The auditory stimulus was a 500 Hz beep (50 dB SPL) without a temporal window due to its short duration, which was played by a loudspeaker located behind the center of the screen. Some visual and auditory stimuli were of higher intensity, the parameters of these stimuli were determined individually (see Intensity-discrimination task). We adjusted the timing of audiovisual stimulus presentation and verified the timing using an oscilloscope (PICOSCOPE 2204A).

### 4.3 Procedure

The experiment consisted of nine sessions, which took place on nine separate days. In each session, participants completed a pre-test, an exposure, and a post-test phase in sequence. The adapter SOA was fixed within a session, but varied across sessions (*±*700, *±*300, *±*200, *±*100, 0 ms). The order of the adapter SOA was randomized across participants, with sessions separated by at least one day. The intensities of the oddball stimuli were determined prior to the experiment for each participant using an intensity-discrimination task to equate the difficulty of detecting oddball stimuli between participants and across modalities.

#### 4.3.1 Pre-test phase

Participants completed a ternary TOJ task during the pre-test phase. Each trial started a fixation cross (0.1–0.2 s, uniform distribution; Fig. 1A), followed by a blank screen (0.4–0.6 s, uniform distribution). Then, an auditory and a visual stimulus (0.033 s) were presented with a variable SOA. There were a total of 15 possible test SOAs (*±*0.5 s and from -0.3 to 0.3 s in steps of 0.05 s), with positive values representing visual lead and negative values representing auditory lead. Following stimulus presentation there was another blank screen (0.4–0.6 s, uniform distribution), and then a response probe appeared on the screen. Participants indicated by button press whether the auditory stimulus occurred before or after the visual stimulus, or the two were simultaneous. There was no time limit for the response, and response feedback was not provided. The inter-trial interval (ITI) was 0.2–0.4 s (uniform distribution). Each test SOA was presented 20 times in pseudo-randomized order, resulting in 300 trials in total, divided into five blocks. Participants usually took around 15 minutes to finish the pre-test phase.

#### 4.3.2 Exposure phase

Participants completed an oddball-detection task during the exposure phase. In each trial, participants were presented with an audiovisual stimulus pair with a fixed SOA (adapter SOA). In 10% of trials, the intensity of either the visual or the auditory component (or both) was greater than in the other trials. Participants were instructed to press the corresponding button as soon as possible to indicate whether there was an auditory oddball, a visual oddball, or both stimuli were oddballs. The task timing (Fig. 1B) was almost identical to the ternary TOJ task, except that there was a response time limit of 1.4 s. Prior to the exposure phase, participants practiced the task for as long as needed to familiarize themselves with the task. During this practice, they were presented with bimodal stimuli with the same adapter SOA used in the exposure phase. There were a total of 250 trials, divided into five blocks. At the end of each block, we presented a performance summary with the hit rate and false alarm rate of each modality. Participants usually took 15 minutes to complete the exposure phase.

#### 4.3.3 Post-test phase

Participants completed the ternary TOJ task as well as the oddball-detection task during the post-test phase. Specifically, each temporal-order judgment was preceded by three top-up (oddball-detection) trials. The adapter SOA in the top-up trials was the same as that in the exposure phase to prevent dissipation of temporal recalibration (Machulla et al., 2012). Both visual and auditory *d*^*′*^ remained consistent from the exposure to post-test phases, indicating similar performance in the top-up trials to performance during the exposure phase (Supplement Figure S10). To facilitate task switching, the ITI between the last top-up trial and the following TOJ trial was longer (with the additional time jittered around 1 s). Additionally, the fixation cross was displayed in red to signal the start of a TOJ trial. As in the pre-test phase, there were 300 TOJ trials (15 test SOAs *×* 20 repetitions) with the addition of 900 top-up trials, grouped into six blocks. At the end of each block, we provided a summary of the oddball-detection performance. Participants usually took around 1 hour to complete the post-test phase.

#### 4.3.4 Intensity-discrimination task

This task was conducted to estimate the just-noticeable-difference (JND) in intensity for a standard visual stimulus with a luminance of 36.1 cd/m^2^ and a standard auditory stimulus with a volume of 40 dB SPL. The task was two-interval, forced choice. The trial started with a fixation (0.1–0.2 s) and a blank screen (0.4–0.6 s). Participants were presented with a standard stimulus (0.033 s) in one randomly selected interval and a comparison stimulus (0.033 s) in the other interval, temporally separated by an inter-stimulus interval (0.6–0.8 s). They indicated which interval contained the brighter/louder stimulus without time constraint. Seven test stimulus levels (luminance range: 5%–195% relative to the standard visual stimulus intensity; volume range: 50%–150% relative to the standard auditory stimulus’ amplitude) were repeated 20 times, resulting in 140 trials for each task. We fit a cumulative Gaussian distribution function to these data and defined the oddball as an auditory or visual stimulus with an intensity judged as more intense than the standard 90% of the time. A higher probability than the standard JND of 75% was selected because the pilot studies showed that the harder oddball detection task became too demanding during the one-hour post-test.

### 4.4 Modeling

In this section, we first outline general assumptions, shared across all candidate models, regarding sensory noise, measurements, and bias. Then, we formalize three process models of recalibration that differ in the implementation of recalibration. In each recalibration model, we also provide a formalization of the ternary TOJ task administered in the pre- and the post-test phases, data from which were used to constrain the model parameters. Finally, we describe how the models were fit to the data.

#### 4.4.1 General modal assumptions regarding sensory noise, measurements and bias

When an audiovisual stimulus pair with a SOA, *s* = *t*_*A*_ − *t*_*V*_, is presented, it triggers auditory and visual signals that are registered in the relevant region of cortex where audiovisual temporal-order comparisons are made. This leads to two internal measurements of the arrival time for each signal in an observer’s brain. These arrival times are subject to noise and thus vary across presentations of the same physical stimulus pair. As in previous work (García-Pérez & Alcalá-Quintana, 2012), we model the probability distribution of the arrival time as shifted exponential distributions (Figure 3A). The arrival time of the auditory signal relative to onset *t*_*A*_ is the sum of the fixed delay of internal signal, *β*_*A*_, and an additional random delay that is exponentially distributed with time constant *τ*_*A*_; analogous for the visual latency (with delay *β*_*V*_ and time constant *τ*_*V*_).

The measured SOA of the audiovisual stimulus pair is modeled as the difference of the arrival times of the two stimuli. Thus, the sensory measurement of SOA, *m*, reflects the sum of three components: the physical SOA, *s*; a fixed latency that is the difference between the auditory and visual fixed delay, *β*_pre_ = *β*_*A*_ −*β*_*V*_ ; and the difference between two exponentially distributed random delays. A negative value of *β*_pre_ indicates faster auditory processing. We assume that the audiovisual fixed latency corresponds to the observer’s default audiovisual temporal bias (Badde, Ley, et al., 2020; Grabot & van Wassenhove, 2017). Thus, we assume that after leaving the experimental room, the default bias is restored and thus consistent across pre-tests.

We model the recalibration process as a shift of the audiovisual temporal bias at the end of every exposure trial *i, β*_*i*_ = *β*_pre_ +Δ_*β,i*_, where *β*_*i*_ is the current audiovisual bias, and Δ_*β,i*_ is the cumulative shift of audiovisual temporal bias. After the 250 exposure trials the updated biases can be expresses as *β*_post_ = *β*_pre_ +Δ_*β*,250_. We also assume that the amounts of auditory and visual latency noise, *τ*_*A*_ and *τ*_*V*_, remain constant across phases and sessions.

Given that both latency distributions are shifted exponential distributions, the probability density function of the sensory measurements of SOA, *m*, given physical SOA, *s*, is a double-exponential function (see derivation in Supplement Section 3; Figure 6A):

**Figure 6:**
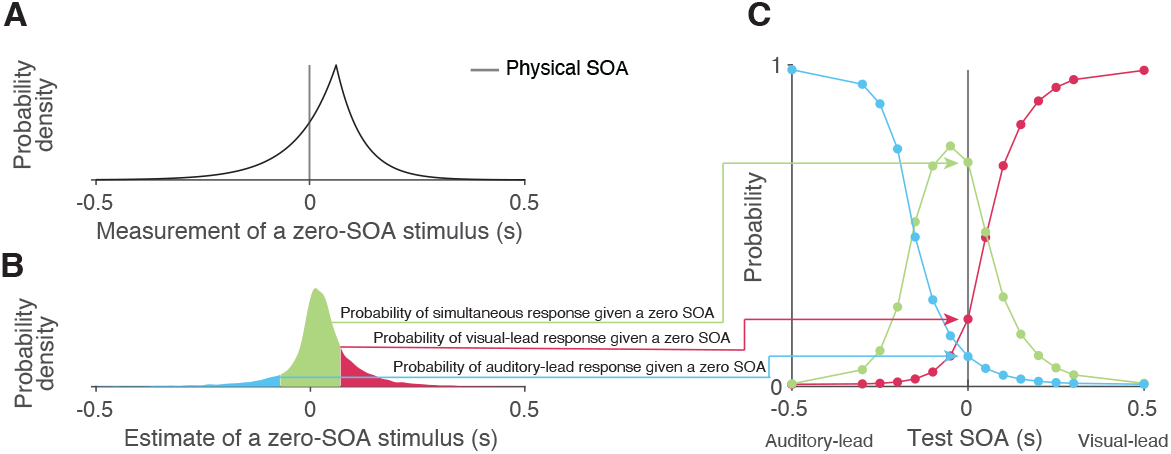
Simulating responses of the TOJ task with a causal-inference perceptual process. (A) An example probability density for the measurement of a zero SOA. (B) The probability density of estimates resulting from a zero-SOA stimulus based on simulation using the causal-inference process. The symmetrical criteria around zero partition the distribution of estimated SOA into three regions, coded by different colors. The area under each segment of the estimate distribution corresponds to the probabilities of the three possible intended responses for a zero SOA. (C) The simulated psychometric function computed by repeatedly calculating the probabilities of the three response types across all test SOAs.

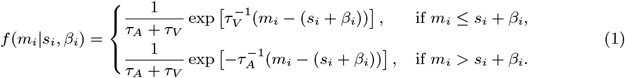

The probability density function of measured SOA peaks at the physical SOA of the stimuli plus the participant’s audiovisual temporal bias, *s*_*i*_ + *β*_*i*_. The left and right spread of this measurement distribution depends on the amount of the latency noise for the visual, *τ*_*V*_, and auditory, *τ*_*A*_, signals. In models with modality-independent arrival-time precision, *τ*_*A*_ = *τ*_*V*_ and the measurement distribution is symmetrical. This symmetrical measurement distribution is often approximated by a Gaussian distribution to fit TOJ responses in previous temporal-recalibration studies (Di Luca et al., 2009; Fujisaki et al., 2004; Harrar & Harris, 2005; Keetels & Vroomen, 2007; Navarra et al., 2005; Tanaka et al., 2011; Vatakis et al., 2007, 2008; Vroomen et al., 2004). Note that we assume the observer has perfect knowledge of the visual and auditory latency noise. Thus, the density of the measurement distribution corresponds to the likelihood function during the inference process when the observer only has the noisy measurement, *m*, and needs to infer the physical SOA, *s*.

#### 4.4.2 The causal-inference model

##### Formalization of recalibration in the exposure phase

The causal-inference model assumes that, at the end of every exposure trial *i*, a discrepancy between the measured SOA, *m*_*i*_, and the final estimate of the stimulus SOA, *ŝ*_*i*_, signals the need for recalibration. The cumulative shift of audiovisual temporal bias Δ_*β,i*_ after exposure trial *i* is,

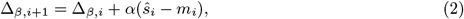

where *α* is the learning rate.

The ideal observer infers intermediate location estimates for two causal scenarios: the auditory and visual stimuli can arise from a single cause (*C* = 1) or two independent causes (*C* = 2). The posterior distribution of the SOA, *s*, conditioned on each causal scenario is computed by multiplying the likelihood function (Eq. 1) with the corresponding prior over SOA. In the case of a common cause (*C* = 1), the prior distribution of the SOA between sound and light is a Gaussian distribution (Magnotti et al., 2013; McGovern et al., 2016), 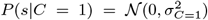. To maintain consistency with previous studies, we used an unbiased prior which assigns the highest probability to a physically synchronous stimulus pair *s* = 0. Similarly, the prior distribution conditioned on separate causes (*C* = 2) is also a Gaussian distribution, 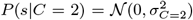, with a much larger spread compared to the common-cause scenario. The intermediate estimates *ŝ*_*C*=1_ conditioned on the commoncause scenario and *ŝ*_*C*=2_ conditioned on separate-cause scenario are the maximum-a-posteriori estimates of conditional posteriors, which are approximated numerically as there is no closed-form solution.

The final estimate of the stimulus SOA, *ŝ*, depends on the posterior probability of each causal scenario. According to Bayes Rule, the posterior probability that an audiovisual stimulus pair with the measured SOA, *m*, shares a common cause is

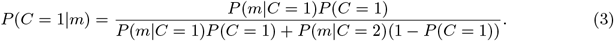

The likelihood of a common source/separate sources for a fixed SOA measurement was approximated by numerically integrating the scenario-specific protoposterior (i.e., the unnormalized posterior),

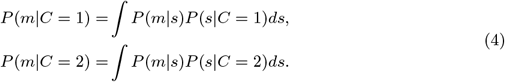

The posterior probability of a common cause additionally depends on the observer’s prior belief of a common cause for auditory and visual stimuli, *P* (*C* = 1) = *p*_common_.

The final estimate of SOA was derived by model averaging, i.e., the average of the scenariospecific SOA estimates, *ŝ*_*C*=1_ and *ŝ*_*C*=2_ each weighted by the posterior probability of the corresponding causal scenario,

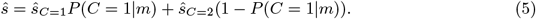

##### Formalization of the ternary TOJ task with a causal-inference perceptual process

In the ternary TOJ task administered in the pre- and post-test phases, the observer is presented with an audiovisual stimulus pair and has to decide whether the auditory stimulus was presented first, the visual stimulus was presented first, or both of them were presented at the same time. The observer makes this perceptual judgment by comparing the final estimate of the SOA, *ŝ*, to two internal criteria (Cary et al., 2024; García-Pérez & Alcalá-Quintana, 2012). We assume that the observer has a symmetric pair of criteria, *±c*, centered on the stimulus SOA corresponding to perceptual simultaneity (*ŝ* = 0). In addition, the observer may lapse or make an error when responding by a lapse rate, *λ*. The probabilities of reporting visual lead, Ψ_*V*_, auditory lead, Ψ_*A*_, or that the two stimuli were simultaneous, Ψ_*S*_, are thus

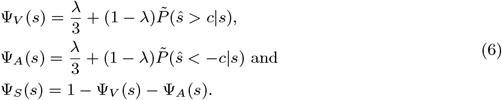

The probability distribution of causal-inference-based SOA estimates *P* (*ŝ*|*s*) has no closed form distribution function and thus was approximated using simulations, resulting in 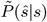. Figure 6 illustrates the process of simulating the psychometric functions, using a zero test SOA as an example. First, we sampled 10,000 SOA measurements from the double-exponential probability distribution corresponding to the test SOA of zero (Figure 6A). Second, for each sampled measurement, we simulated the process by which the observer carries out causal inference and by doing so produced an estimate of the stimulus SOA, while keeping the causal-inference model parameters fixed. This process resulted in a Monte-Carlo approximation of the probability density distribution of the causal-inference-based SOA estimates (Figure 6B). Third, we calculated the probability of the three types of responses (Eq. 6) for this specific test SOA. This process was repeated for each test SOA to generate three psychometric functions (Figure 6C).

#### 4.4.3 The asynchrony-contingent model

In the asynchrony-contingent model, the observer measures the audiovisual SOA, *s*, by comparing the arrival latency of the auditory and visual signals. The observer uses the likelihood that the audiovisual stimuli occurred simultaneously *P* (*m*|SOA = 0) to update the temporal bias during recalibration, instead of performing causal inference. We again assume that the observer has perfect knowledge about the variability and fixed delays of the arrival times and thus assume the likelihood corresponds to the measurement distribution (Eq. 1). The observer uses this probability of simultaneity to scale the update rate of the audiovisual bias,

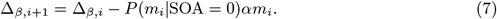

We assume the observer’s estimate of the stimulus SOA, *ŝ*, is identical to the measured SOA, *m*. Thus, from the experimenter’s perspective, the probability of the three different responses in the TOJ task can be obtained by replacing the SOA estimate, *ŝ*, with the SOA measurement, *m*, in Eq. 6). As we know the probability distribution of *m*, the psychometric functions have a closed form (García-Pérez & Alcalá-Quintana, 2012).

#### 4.4.4 The asynchrony-correction model

In the asynchrony-correction model, the observer begins by evaluating if the sensory measurement of SOA, *m*, falls outside the criterion range for reporting that the two stimuli were presented simultaneously ±*c*. If the measurement does exceed this criterion, the observer adjusts the audiovisual bias by shifting it against the measurement, i.e., shifting it so that the measured SOA of a pair would be closer to zero and is more likely to perceived as simultaneous. This adjustment is proportional to the sensory measurement of the SOA, *m*, at a fixed rate determined by the learning rate *α*. The update rule of the audiovisual bias in trial *i* is thus

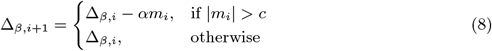

The derivation of the psychometric functions is identical to the asynchrony-contingent model.

#### 4.4.5 Model fitting

##### Model log-likelihood

The model was fitted by optimizing the lower bound on the marginal log-likelihood. We fit the model to the ternary TOJ data collected during the pre- and post-test phases of all sessions together. We did not collect temporal-order judgments in the exposure phase. But, to model the post-test data, we need to estimate the distribution of shifts of audiovisual bias resulting from the exposure phase (Δ_*β*,250_). We do this using Monte Carlo simulation of the 250 exposure trials to estimate the probability distribution of the cumulative shifts.

The set of model parameters Θ is listed in Table 1. There are *J* sessions, each including *K* trials in the pre-test phase and *K* trials in the post-test phase. We denote the full dataset of pre-test data as *X*_pre_ and for the post-test data as *X*_post_. We fit the pre- and post-test data jointly by summing their log-likelihood, log *p*(*X*|*M*, Θ) = log *p*(*X*_pre_|*M*, Θ) + log *p*(*X*_post_|*M*, Θ).

**Table 1:** Model parameters. Check marks signify that the parameter is used for determining the likelihood of the data from the temporal-order judgment task in the pre- and post-test phase and/or for the Monte Carlo simulation of recalibration in the exposure phase.

| Notation | Specification | Temporal-order-judgement task | Recalibration in the exposure phase |
| --- | --- | --- | --- |
| $\beta_{\text{pre}}$ | The fixed relative delay between visual and auditory processing, i.e., the audio-visual bias prior to the exposure phase | ✓ | ✓ |
| $\tau_A$ | Amount of auditory latency noise, the exponential time constant of the auditory detection-latency distribution | ✓ | ✓ |
| $\tau_V$ | Amount of visual latency noise, the exponential time constant of the visual detection-latency distribution | ✓ | ✓ |
| $\sigma_{C=1}$ | The spread of the Gaussian prior for the common-cause scenario | ✓ | ✓ |
| $\sigma_{C=2}$ | The spread of the Gaussian prior for the separate-causes scenario | ✓ | ✓ |
| $p_{\text{common}}$ | The prior probability of a common cause | ✓ | ✓ |
| $c$ | Simultaneity criterion | ✓ | |
| $\lambda$ | Lapse rate | ✓ | |
| $\alpha$ | Learning rate for shifting audiovisual bias | | ✓ |

In a given trial, the observer responded either auditory-first (A), visual-first (V), or simultaneous (S). We denote a single response using indicator variables that are equal to 1 if that was the response in that trial and 0 otherwise. These variables for trial *k* in session *j* are 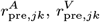 and 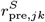 for the pre-test trials, and 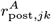, etc., for the post-test trials.

The log-likelihood of all pre-test responses *X*_pre_ given the model parameters is

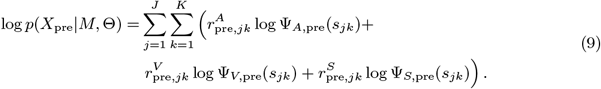

The psychometric functions for the pre-test (e.g., Ψ_*A*,pre_) are defined in Eq. 6, and are the same across all sessions as we assumed that the audiovisual bias *β*_pre_ was the same before recalibration in every session.

The log-likelihood of responses in the post-test depends on the audiovisual bias after recalibration *β*_post,*j*_ = *β*_pre_ + Δ_*β*,250,*j*_ for session *j*. To determine the log-likelihood of the post-test data requires us to integrate out the unknown value of the cumulative shift Δ_*β*,250,*j*_. We approximated this integral in two steps based on our previous work (Hong et al., 2021). First, we simulated the 250 exposure trials 1000 times for a given set of parameters Θ and session *j*. This resulted in 1,000 values of Δ_*β*,250,*j*_. The distribution of these values was well fit by a Gaussian whose parameters were determined by the empirical mean and standard deviation of the sample distribution, resulting in the distribution 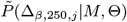. Second, we approximated the integral of the log-likelihood of the data over possible values of Δ_*β*,250,*j*_ by numerical integration. We discretized the approximated distribution 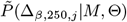 into 100 equally spaced bins centered on values Δ_*β*,250,*j*_(*n*) (*n* = 1, …, 100). The range of the bins was triple the range of the values from the Monte Carlo sample, so that the lower bound was 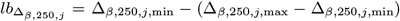 and the upper bound was 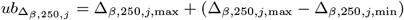.

The log-likelihood of the post-test data was approximated as

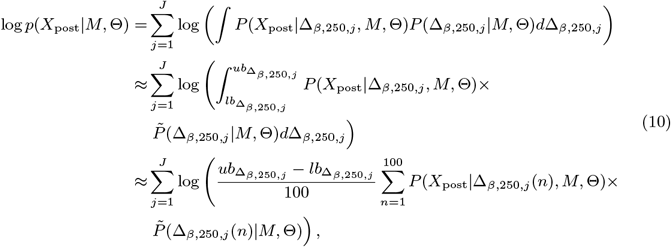

where

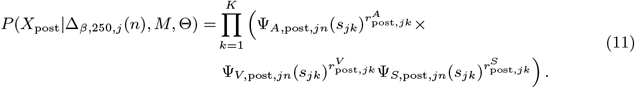

The psychometric functions in the post-test (e.g., Ψ_*A*,post,*jn*_) differed across sessions and bins because the simulated audiovisual bias after the exposure phase *β*_post,*j*_ depends on the adapter SOA fixed in session *j* and the simulation bin *n*.

##### Parameter estimation and model comparison

We approximated the lower bounds to the model evidence (i.e., the marginal likelihood) of each model for each participant’s data using Variational Bayesian Monte Carlo (Acerbi, 2018, 2020). We set the prior distribution of parameters based on the results of maximum likelihood estimation using Bayesian Adaptive Direct Search to ensure that the parameter ranges were plausible (Acerbi & Ma, 2017). We repeated each search 20 times with a different and random starting point to address the possibility of reporting a local minimum. For each model, the fit with the maximum lower bounds of the model evidence across the repeated searches was chosen for the maximum model evidence and best parameter estimates.

We then conducted a Bayesian model comparison based on model evidence. The model with the strongest evidence was considered the best-fitting model (MacKay, 2003). To quantify the support of model selection, we computed the Bayes factor, the ratio of the model evidence between each model and the asynchrony-correction, modality-independent-precision model, which had the weakest model evidence. To compare any two models, one can simply calculate the difference in their log Bayes factors as both are relative to the same weakest model.

##### Model recovery and parameter recovery

We conducted a model-recovery analysis for the six models and confirmed that they are identifiable (Supplement Section 11). In addition, we considered an alternative causal-inference model in which the bias update is proportional to the posterior probability of a common cause, instead of driven by the percept. A separate model recovery analysis on variations of the causal-inference model was unable to distinguish between them (Supplement Section 12). For the causal-inference, modality-specific-precision model, we also carried out a parameter recovery analysis and confirmed that all the parameters are recoverable (Supplement Section 13).

## 5 Declarations

### 5.1 Data and code availability

All data and code are available via the Open Science Framework (https://osf.io/8s7qv/).

## 5.2 Acknowledgments

We thank the NYU High-Performance Computing (NYU HPC) for providing computational resources and support. We thank the anonymous reviewers’ advice that helped us improve the manuscript.

## References

Acerbi, L. (2018). Variational Bayesian Monte Carlo. arXiv [stat.ML].

Acerbi, L. (2020). Variational Bayesian Monte Carlo with noisy likelihoods. Advances in Neural Information Processing Systems, 33, 8211–8222.

Acerbi, L., & Ma, W. J. (2017). Practical bayesian optimization for model fitting with bayesian adaptive direct search. Proceedings of the 31st International Conference on Neural Information Processing Systems, 1834–1844.

Badde, S., Landy, M. S., & Adams, W. J. (2023). Multisensory causal inference is feature-specific, not object-based. Philos. Trans. R. Soc. Lond. B Biol. Sci., 378 (1886), 20220345.

Badde, S., Ley, P., Rajendran, S. S., Shareef, I., Kekunnaya, R., & Röder, B. (2020). Sensory experience during early sensitive periods shapes cross-modal temporal biases. Elife, 9, Article e61238.

Badde, S., Navarro, K. T., & Landy, M. S. (2020). Modality-specific attention attenuates visual-tactile integration and recalibration effects by reducing prior expectations of a common source for vision and touch. Cognition, 197, Article 104170.

Beierholm, U. R., Quartz, S. R., & Shams, L. (2009). Bayesian priors are encoded independently from likelihoods in human multisensory perception. J. Vis., 9 (5), Article 23.

Berniker, M., & Kording, K. (2008). Estimating the sources of motor errors for adaptation and generalization. Nat. Neurosci., 11 (12), 1454–1461.

Cai, M., Stetson, C., & Eagleman, D. M. (2012). A neural model for temporal order judgments and their active recalibration: A common mechanism for space and time? Front. Psychol., 3, Article 470.

Cary, E., Lahdesmaki, I., & Badde, S. (2024). Audiovisual simultaneity windows reflect temporal sensory uncertainty. Psychon. Bull. Rev.

Di Luca, M., Machulla, T.-K., & Ernst, M. O. (2009). Recalibration of multisensory simultaneity: Cross-modal transfer coincides with a change in perceptual latency. J. Vis., 9 (12), Article 7.

Fain, G. L. (2019). Sensory transduction (2nd ed.). Oxford University Press.

Fujisaki, W., Shimojo, S., Kashino, M., & Nishida, S. (2004). Recalibration of audio-visual simultaneity. Nat. Neurosci., 7 (7), 773–778.

García-Pérez, M. A., & Alcalá-Quintana, R. (2012). On the discrepant results in synchrony judgment and temporal-order judgment tasks: A quantitative model. Psychon. Bull. Rev., 19 (5), 820–846.

Grabot, L., & van Wassenhove, V. (2017). Time order as psychological bias. Psychol. Sci., 28 (5), 670–678.

Hanson, J. V. M., Heron, J., & Whitaker, D. (2008). Recalibration of perceived time across sensory modalities. Exp. Brain Res., 185 (2), 347–352.

Harrar, V., & Harris, L. R. (2005). Simultaneity constancy: Detecting events with touch and vision. Exp. Brain Res., 166 (3-4), 465–473.

Harrar, V., & Harris, L. R. (2008). The effect of exposure to asynchronous audio, visual, and tactile stimulus combinations on the perception of simultaneity. Exp. Brain Res., 186 (4), 517–524.

Heron, J., Roach, N. W., Hanson, J. V. M., McGraw, P. V., & Whitaker, D. (2012). Audiovisual time perception is spatially specific. Exp. Brain Res., 218 (3), 477–485.

Heron, J., Whitaker, D., McGraw, P. V., & Horoshenkov, K. V. (2007). Adaptation minimizes distance-related audiovisual delays. J. Vis., 7 (13), Article 5.

Hirsh, I. J., & Sherrick, C. E., Jr. (1961). Perceived order in different sense modalities. J. Exp. Psychol., 62 (5), 423–432.

Hong, F. (2023, May). The role of causal inference in multisensory integration and recalibration [Doctoral dissertation, New York University].

Hong, F., Badde, S., & Landy, M. S. (2021). Causal inference regulates audiovisual spatial recalibration via its influence on audiovisual perception. PLoS Comput. Biol., 17 (11), Article e1008877.

Hsiao, A., Lee-Miller, T., & Block, H. J. (2022). Conscious awareness of a visuo-proprioceptive mismatch: Effect on cross-sensory recalibration. Front. Neurosci., 16, Article 958513.

Keetels, M., & Vroomen, J. (2007). No effect of auditory-visual spatial disparity on temporal recalibration. Exp. Brain Res., 182 (4), 559–565.

King, A. J. (2005). Multisensory integration: Strategies for synchronization. Curr. Biol., 15 (9), R339–41.

Körding, K. P., Beierholm, U., Ma, W. J., Quartz, S., Tenenbaum, J. B., & Shams, L. (2007). Causal inference in multisensory perception. PLoS One, 2 (9), Article e943.

Machulla, T.-K., Di Luca, M., Froehlich, E., & Ernst, M. O. (2012). Multisensory simultaneity recalibration: Storage of the aftereffect in the absence of counterevidence. Exp. Brain Res., 217 (1), 89–97.

MacKay, D. J. C. (2003). Information theory, inference and learning algorithms. Cambridge University Press.

Magnotti, J. F., Ma, W. J., & Beauchamp, M. S. (2013). Causal inference of asynchronous audiovisual speech. Front. Psychol., 4, Article 798.

Maij, F., Brenner, E., & Smeets, J. B. J. (2009). Temporal information can influence spatial localization. J. Neurophysiol., 102 (1), 490–495.

McGovern, D. P., Roudaia, E., Newell, F. N., & Roach, N. W. (2016). Perceptual learning shapes multisensory causal inference via two distinct mechanisms. Sci. Rep., 6, Article 24673.

Navarra, J., Hartcher-O’Brien, J., Piazza, E., & Spence, C. (2009). Adaptation to audiovisual asynchrony modulates the speeded detection of sound. Proc. Natl. Acad. Sci. U. S. A., 106 (23), 9169–9173.

Navarra, J., Vatakis, A., Zampini, M., Soto-Faraco, S., Humphreys, W., & Spence, C. (2005). Exposure to asynchronous audiovisual speech extends the temporal window for audiovisual integration. Brain Res. Cogn. Brain Res., 25 (2), 499–507.

O’Donohue, M., Lacherez, P., & Yamamoto, N. (2022). Musical training refines audiovisual integration but does not influence temporal recalibration. Sci. Rep., 12, Article 15292.

Pöppel, E. (1988). Mindworks: Time and conscious experience. Harcourt Brace Jovanovich.

Roach, N. W., Heron, J., Whitaker, D., & McGraw, P. V. (2011). Asynchrony adaptation reveals neural population code for audio-visual timing. Proc. Biol. Sci., 278 (1710), 1314–1322.

Rohde, M., & Ernst, M. O. (2012). To lead and to lag - forward and backward recalibration of perceived visuo-motor simultaneity. Front. Psychol., 3, Article 599.

Rohde, M., Greiner, L., & Ernst, M. O. (2014). Asymmetries in visuomotor recalibration of time perception: Does causal binding distort the window of integration? Acta Psychol., 147, 127–135.

Rohe, T., & Noppeney, U. (2015). Sensory reliability shapes perceptual inference via two mechanisms. J. Vis., 15 (5), Article 22.

Roseboom, W., & Arnold, D. H. (2011). Twice upon a time: Multiple concurrent temporal recalibrations of audiovisual speech. Psychol. Sci., 22 (7), 872–877.

Roseboom, W., Kawabe, T., & Nishida, S. (2013). Audio-Visual temporal recalibration can be constrained by content cues regardless of spatial overlap. Front. Psychol., 4, Article 189.

Roseboom, W., Linares, D., & Nishida, S. (2015). Sensory adaptation for timing perception. Proc. Biol. Sci., 282 (1805), Article 20142833.

Sato, Y. (2021). Comparing bayesian models for simultaneity judgement with different causal assumptions. J. Math. Psychol., 102 (102521), 102521.

Sato, Y., & Aihara, K. (2011). A bayesian model of sensory adaptation. PLoS One, 6 (4), Article e19377.

Sato, Y., Toyoizumi, T., & Aihara, K. (2007). Bayesian inference explains perception of unity and ventriloquism aftereffect: Identification of common sources of audiovisual stimuli. Neural Comput., 19 (12), 3335–3355.

Shams, L., & Beierholm, U. R. (2010). Causal inference in perception. Trends Cogn. Sci., 14 (9), 425–432.

Spence, C., & Squire, S. (2003). Multisensory integration: Maintaining the perception of synchrony. Curr. Biol., 13 (13), R519–21.

Sternberg, S., & Knoll, R. L. (1973). The perception of temporal order: Fundamental issues and a general model.

Tanaka, A., Asakawa, K., & Imai, H. (2011). The change in perceptual synchrony between auditory and visual speech after exposure to asynchronous speech. Neuroreport, 22 (14), 684–688.

Van der Burg, E., Alais, D., & Cass, J. (2013). Rapid recalibration to audiovisual asynchrony. J. Neurosci., 33 (37), 14633–14637.

van Beers, R. J., Wolpert, D. M., & Haggard, P. (2002). When feeling is more important than seeing in sensorimotor adaptation. Curr. Biol., 12 (10), 834–837.

Vatakis, A., Navarra, J., Soto-Faraco, S., & Spence, C. (2007). Temporal recalibration during asynchronous audiovisual speech perception. Exp. Brain Res., 181 (1), 173–181.

Vatakis, A., Navarra, J., Soto-Faraco, S., & Spence, C. (2008). Audiovisual temporal adaptation of speech: Temporal order versus simultaneity judgments. Exp. Brain Res., 185 (3), 521–529.

Vroomen, J., & de Gelder, B. (2004). Temporal ventriloquism: Sound modulates the flash-lag effect. J. Exp. Psychol. Hum. Percept. Perform., 30 (3), 513–518.

Vroomen, J., & Keetels, M. (2010). Perception of intersensory synchrony: A tutorial review. Atten. Percept. Psychophys., 72 (4), 871–884.

Vroomen, J., Keetels, M., de Gelder, B., & Bertelson, P. (2004). Recalibration of temporal order perception by exposure to audio-visual asynchrony. Brain Res. Cogn. Brain Res., 22 (1), 32–35.

Wei, K., & Körding, K. (2009). Relevance of error: What drives motor adaptation? J. Neurophysiol., 101 (2), 655–664.

Wozny, D. R., Beierholm, U. R., & Shams, L. (2010). Probability matching as a computational strategy used in perception. PLoS Comput. Biol., 6 (8), Article e1000871.

Yarrow, K., Jahn, N., Durant, S., & Arnold, D. H. (2011). Shifts of criteria or neural timing? the assumptions underlying timing perception studies. Conscious. Cogn., 20 (4), 1518–1531.

Yarrow, K., Kohl, C., Segasby, T., Kaur Bansal, R., Rowe, P., & Arnold, D. H. (2022). Neural-latency noise places limits on human sensitivity to the timing of events. Cognition, 222, Article 105012.

Yarrow, K., Minaei, S., & Arnold, D. H. (2015). A model-based comparison of three theories of audiovisual temporal recalibration. Cogn. Psychol., 83, 54–76.

Yarrow, K., Roseboom, W., & Arnold, D. H. (2011). Spatial grouping resolves ambiguity to drive temporal recalibration. J. Exp. Psychol. Hum. Percept. Perform., 37 (5), 1657–1661.

Yuan, X., Li, B., Bi, C., Yin, H., & Huang, X. (2012). Audiovisual temporal recalibration: Space-based versus context-based. Perception, 41 (10), 1218–1233.

