## Supplementary Materials for "Precision-based causal inference modulates audiovisual temporal recalibration"

**2 Individual-level empirical data of the recalibration effect 5**

**3 Derivation of the double-exponential distribution of arrival-latency difference 6**

**4 Individual-level model comparison of the recalibration models 8**

**5 Individual-level model predictions of recalibration effects 9**

**7 Simulation of the causal-inference model 16**

**8 Simulation of a population-code recalibration model 17**

**9 Exclusion of an outlier participant 19**

**10 Performance in the oddball-detection task 20**

**11 Model recovery of recalibration models 21**

**12 Model recovery of variants of the causal-inference recalibration models 22**

**13 Parameter recovery of the causal-inference, modality-specific-precision model 23**

### 1 Comparing atheoretical models of audiovisual temporal recalibration

Figure 2 in the main text shows raw psychometric data with fit psychometric functions (panel A) and estimates of point of subjective simultaneity (PSS) shift across sessions (panel B). These estimates are from fitting an *atheoretical* model that estimates PSS shift without reference to an underlying model of how recalibration comes about. Rather, it aims to show the pattern of recalibration from the data alone. Here, we describe how those estimates were computed and compare four different models of recalibration for this purpose. These four models vary based on two factors: the use of either a Gaussian or an exponential measurement distribution, and the assumption of either a bias shift or a criterion shift. Next, we describe these models followed by a model comparison.

#### 1.1 Gaussian vs. exponential measurement distribution

Models of temporal-order judgments (TOJs) typically assume independent sensory channels for each modality, with the measurement error of arrival latency for each channel being either Gaussian (Schneider & Bavelier, 2003; Sternberg & Knoll, 1973) or exponentially distributed (García-Pérez & Alcalá-Quintana, 2012; Petrini et al., 2020).

If arrival latencies are corrupted by Gaussian-distributed noise, the measurements of difference between auditory and visual arrival latencies $m$ also have a Gaussian distribution,

$$P(m \mid s) = \mathcal{N}(s + \beta,\, \sigma_m^2), \tag{S1}$$

where $s$ is the physical stimulus-onset asynchrony (SOA), $\beta$ is the audiovisual bias, and $\sigma_m^2$ is the sum of the variabilities of auditory and visual latency. This model has a conceptual problem: the continuous Gaussian distribution allows for a nonzero probability of negative arrival latencies, but the auditory or visual signal cannot be registered in the brain before the physical stimulus occurs.

The exponential distribution of arrival latencies avoids this problem. As in the main text, the measurement of audiovisual latency $m$ has a double-exponential distribution (Eq. 1). For both models, the order judgment is then made by comparing the arrival-latency difference and decision criteria.

#### 1.2 Bias-shift vs. criterion-shift model

Audiovisual temporal recalibration can result from a change in the audiovisual bias or decision criteria (Yarrow et al., 2011, 2023). The asymmetry observed in audiovisual temporal recalibration can also be modeled by allowing for asymmetric decision criteria by expanding the simultaneity window on one side (O'Donohue et al., 2022; Rohde et al., 2014; Yarrow et al., 2011). For each model, we assume that either the bias or criterion parameter changes after the recalibration phase, and is restored to a default value before the pre-test of a subsequent session. We fit the TOJ responses jointly across pre- and post-tests and all nine sessions. These TOJ models include a bias or a criterion shift as a session-dependent free parameter and keep the other parameters constant across sessions.

##### 1.2.1 Specification of the bias-shift model

In the decision stage, the observer compares the measured SOA with a simultaneity temporal window bounded by symmetric criteria $\pm c$. Including a lapse rate $\lambda$, the psychometric functions are:

$$\begin{aligned}
\Psi_V(s) &= \frac{\lambda}{3} + (1-\lambda)P(m > c \mid s),\\
\Psi_A(s) &= \frac{\lambda}{3} + (1-\lambda)P(m < -c \mid s),\\
\Psi_S(s) &= 1 - \Psi_V(s) - \Psi_A(s).
\end{aligned} \tag{S2}$$

where $P(m \mid s)$ is a Gaussian or double-exponential distribution.

In the pre-test, the parameters and the resulting predicted psychometric functions are the same across sessions. In the post-test, there is a free parameter for the shift of audiovisual bias $\beta$ after exposure to the adapter SOA within that session. Therefore, there are 9 free bias parameters, $\beta_{\text{post},j}$, one for each of the 9 sessions $j$. In sum, the parameter set $\Theta$ includes either 14 free parameters, $\Theta = \{\tau_A, \tau_V, c, \lambda, \beta_{\text{pre}}, \{\beta_{\text{post},j}\}\}$, using a double-exponential distribution, or 13 for the Gaussian measurement distribution, $\Theta = \{\sigma_m, c, \lambda, \beta_{\text{pre}}, \{\beta_{\text{post},j}\}\}$.

##### 1.2.2 Specification of the criterion-shift model

In the criterion-shift model, the simultaneity window is defined by a lower criterion $c_l$ and an upper criterion $c_u$, which can be asymmetric. The measurement distribution is compared with the simultaneity temporal window to obtain the psychometric functions for the three responses. The psychometric functions are the same as Eq. S2 except that we allow for an expansion of the simultaneity window in the post-test phase, thus replacing $c$ with $c_h$ in the definition of $\Psi_V$ and $-c$ with $-c_l$ in the definition of $\Psi_A$.

In the exposure phase, if the adapter SOA is inside the simultaneity window, no recalibration occurs. On the other hand, if it is outside the simultaneity window, recalibration occurs by shifting outward the criterion that is closer to the adapter SOA, that is, expanding the range considered simultaneous on that side. The other side of the criterion remains unchanged. We assume that in a given session *only* the criterion on the same side as the adapter is ever changed. Thus, each session is characterized by a free parameter for how much that criterion is shifted ($\Delta_{c,j}$ for session $j$), so that either $c_h$ is updated to $c_h + \Delta_{c,j}$ or $c_l$ is updated to $c_l - \Delta_{c,j}$. All pre-test sessions share all parameters (and predicted psychometric functions). Taken together, the parameter set $\Theta$ includes either 14 free paramters, $\Theta = \{\tau_A, \tau_V, \lambda, c_l, c_u, \{\Delta_{c,j}\}\}$, for the double-exponential model, or 13 free parameters for the Gaussian model: $\Theta = \{\sigma_m, \lambda, c_l, c_u, \{\Delta_{c,j}\}\}$.

#### 1.3 Model log-likelihood

We fit each model $M$ separately using the same Variational Bayesian Monte Carlo procedure. The TOJ data from the pre- and post-test phases of all sessions were fit jointly. Note that each model does not simulate recalibration, but fits session-dependent free parameters instead.

The parameter set $\Theta$ is specified above for each model, and $X$ is the data set containing each type of response from $I$ phases, $J$ sessions, and $K$ trials ($I = \{\text{pre}, \text{post}\}, J = 9, K = 300$). We maximized the log-likelihood of model parameters given the data,

$$\begin{aligned} \log p(X|M,\Theta) = \sum_{i \in \{\text{pre},\text{post}\}} \sum_{j=1}^{J} \sum_{k=1}^{K} \Big( & r_{ijk}^{A} \log \Psi_{A,ij}(s_{ijk}) + \\ & r_{ijk}^{V} \log \Psi_{V,ij}(s_{ijk}) + \\ & r_{ijk}^{S} \log \Psi_{S,ij}(s_{ijk}) \Big), \end{aligned} \tag{S3}$$

where the $r_{ijk}$ are indicator variables that are equal to one when the corresponding key was pressed, and zero otherwise. The psychometric functions (e.g., $\Psi_{A,ij}$) for the pre-test are the same across sessions and differ for the post-test across sessions due to recalibration.

#### 1.4 Model comparison

Model comparison based on model evidence found that the bias-shift model with exponential measurement distribution captured the data of most participants. We continued to use this model as the atheoretical model as the baseline to compare with other recalibration models in the main text.

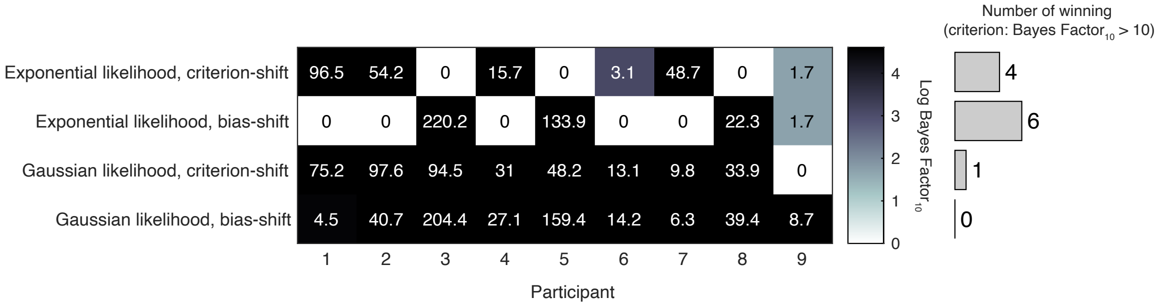

Figure S1: Model comparison results of the atheoretical models. The model with the largest log model evidence is determined to be the best-fitting model for each participant. The table displays the log Bayes Factor of each model relative to the best-fitting model for each participant. Thus, the best-fitting model has a relative log Bayes Factor of zero. Larger values for other models indicate stronger support for the best-fitting model. The vertical histogram shows the number of participants for which each model was the best fit, based on the criterion that a model is rejected if its relative log Bayes Factor is greater than log(10), which is 2.3.

#### 2 Individual-level empirical data of the recalibration effect

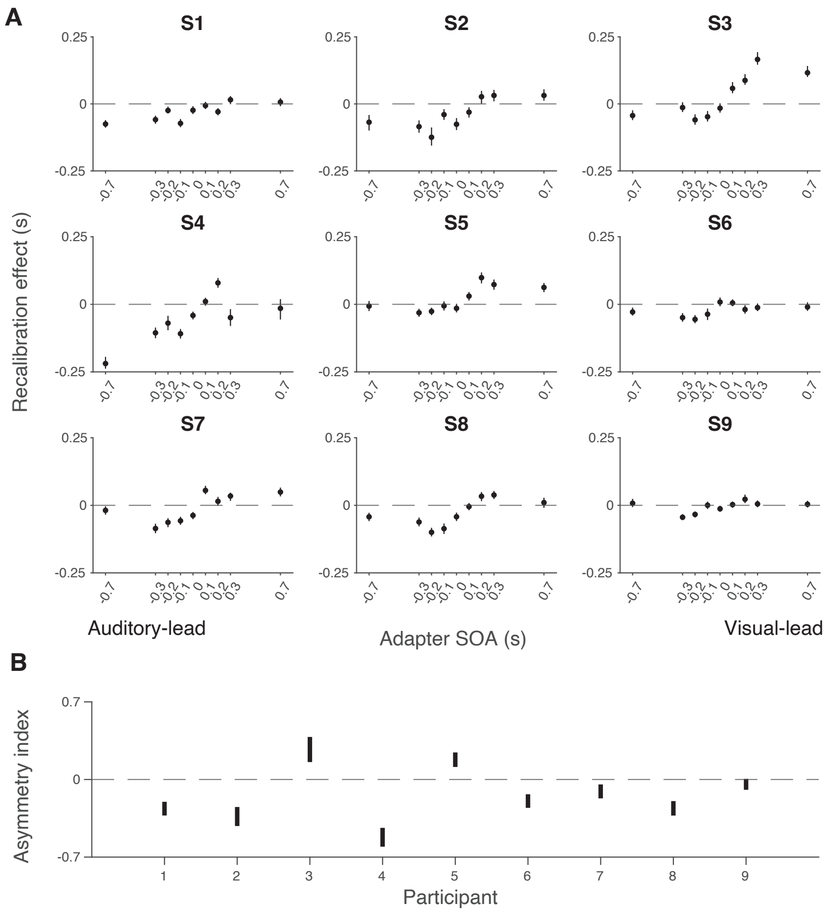

Figure S2: (A) Individual recalibration effect estimated from the atheoretical model that assumes a bias shift and double-exponential measurement distribution. Error bars: 95% confidence interval of recalibration amount from 1000 bootstrapped datasets for that participant. (B) The 95% confidence interval of the asymmetry index (i.e., summed recalibration amount across adapter SOAs) from 1000 bootstrapped datasets for each participant. The confidence interval for all except the last participant excluded zero (CI of S9: [-93.04, 5.85]), suggesting a general asymmetry in recalibration.

### 3 Derivation of the double-exponential distribution of arrival-latency difference

In this section, we derive the asymmetric double-exponential distribution of measured SOA that results from exponential arrival-time distributions (Eq. 1 in the main text). The stimulus onsets occur at times $s_A$ and $s_V$. Each modality has a fixed delay until detection ($\beta_A$ and $\beta_V$). The probability distribution of further delays is an exponential distribution with time constant $\tau_A$ or $\tau_V$. Thus, the arrival time for auditory detection is $T_A = s_A + \beta_A + D_A$, where the random delay $D_A$ is distributed as $f_{D_A}(d_A) = \frac{e^{d_A/\tau_A}}{\tau_A}$, and similarly for the visual stimulus. We are interested in measurements of SOA, that is, the distribution of $D = T_A - T_V = (s_A + \beta_A) - (s_V + \beta_V) + D_A - D_V$. We now derive the distribution for the random variable $D' = D_A - D_V$.

We assume that visual and auditory delays ($D_A$ and $D_V$) are independent. Thus, to compute the density $f_{D'}(d)$ for a particular relative delay $d$, we integrate over all pairs of delays $d_V$ and $d_A = d_V + d$ that correspond to that relative delay such that both individual delays are possible (i.e., both are non-negative). We treat the cases $d \geq 0$ and $d \leq 0$ separately.

**Case 1**: $d \geq 0$

$$f_D(d) = \int_0^\infty f_{D_V}(d_V)\, f_{D_A}(d_V + d)\, dd_V.$$

Here, the lower limit on the integral is zero so that both delays are non-negative. Substituting the PDFs:

$$\begin{aligned} f_D(d) &= \int_0^\infty \tau_V^{-1} e^{-d_V/\tau_V} \cdot \tau_A^{-1} e^{-(d_V+d)/\tau_A}\, dd_V \\ &= \tau_V^{-1}\tau_A^{-1} e^{-d/\tau_A} \int_0^\infty e^{-d_V/\tau_V - d_V/\tau_A}\, dd_V \\ &= \tau_V^{-1}\tau_A^{-1} e^{-d/\tau_A} \int_0^\infty e^{-(\tau_V+\tau_A)d_V/\tau_V\tau_A}\, dd_V \\ &= \tau_V^{-1}\tau_A^{-1} e^{-d/\tau_A} (\tau_V\tau_A)/(\tau_V + \tau_A) \\ &= \frac{e^{-d/\tau_A}}{\tau_V + \tau_A}. \end{aligned} \tag{S4}$$

**Case 2**: $d \leq 0$

$$f_D(d) = \int_{-d}^\infty f_{D'_V}(d_V)\, f_{D'_A}(d_V + d)\, dd_V.$$

Here, the lower limit on the integral is $-d$ (which is a positive value) so that both delays are non-negative. Substituting the PDFs:

$$
\begin{aligned}
f_D(d) &= \int_{-d}^{\infty} \tau_V^{-1} e^{-d_V/\tau_V} \cdot \tau_A^{-1} e^{-(d_V+d)/\tau_A}\, dd_V \\
&= \tau_V^{-1}\tau_A^{-1} e^{-d/\tau_A} \int_{-d}^{\infty} e^{-d_V/\tau_V - d_V/\tau_A}\, dd_V \\
&= \tau_V^{-1}\tau_A^{-1} e^{-d/\tau_A} \int_{-d}^{\infty} e^{-(\tau_V+\tau_A)d_V/\tau_V\tau_A}\, dd_V \\
&= \tau_V^{-1}\tau_A^{-1} e^{-d/\tau_A} (\tau_V\tau_A)/(\tau_V+\tau_A) e^{(\tau_V+\tau_A)d/\tau_V\tau_A} \\
&= e^{-d/\tau_A} e^{(1/\tau_V+1/\tau_A)d}/(\tau_V+\tau_A) \\
&= \frac{e^{d/\tau_V}}{\tau_V+\tau_A}.
\end{aligned} \tag{S5}
$$

Combining the two cases, the PDF of $D$ is:

$$
f_D(d) = \begin{cases} \dfrac{e^{d/\tau_V}}{\tau_V+\tau_A}, & \text{if } d \le 0, \\ \dfrac{e^{-d/\tau_A}}{\tau_V+\tau_A}, & \text{if } d \ge 0. \end{cases} \tag{S6}
$$

The difference of the random delays follows a double exponential distribution centered at zero, with different decay rates on either side of zero determined by $\tau_A$ and $\tau_V$. Adding in the fixed relative delays $((s_A + \beta_A) - (s_V + \beta_V))$ we arrive at Eq. 1 in the main text.

### 4 Individual-level model comparison of the recalibration models

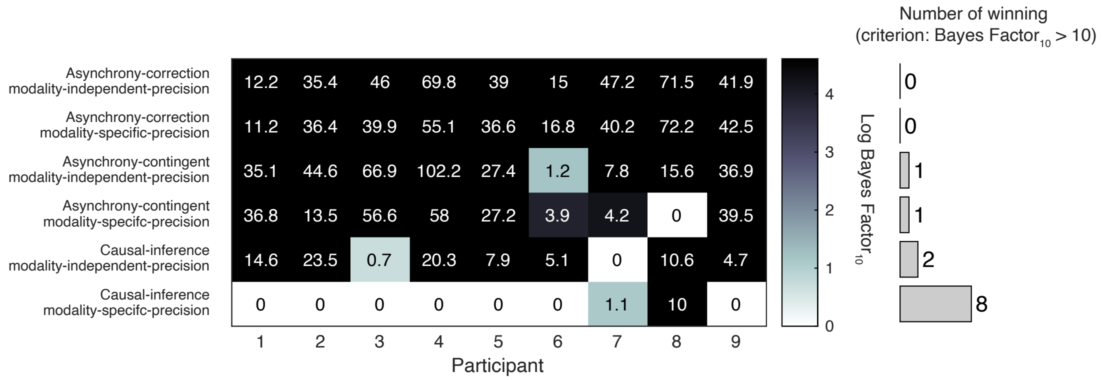

Figure S3: Model comparison results of the recalibration models. The table displays log Bayes Factor relative to the best-fitting model for each participant. Thus, the best-fitting model has a relative log Bayes Factor of 0. Larger values for other models indicate stronger support for the best-fitting model. The vertical histogram shows the number of participants for which each model was the best fit, based on the criterion that a model is rejected if the log Bayes Factor relative to the best-fitting model is greater than log(10), which is 2.3.

### 5 Individual-level model predictions of recalibration effects

#### 5.1 Causal-inference model

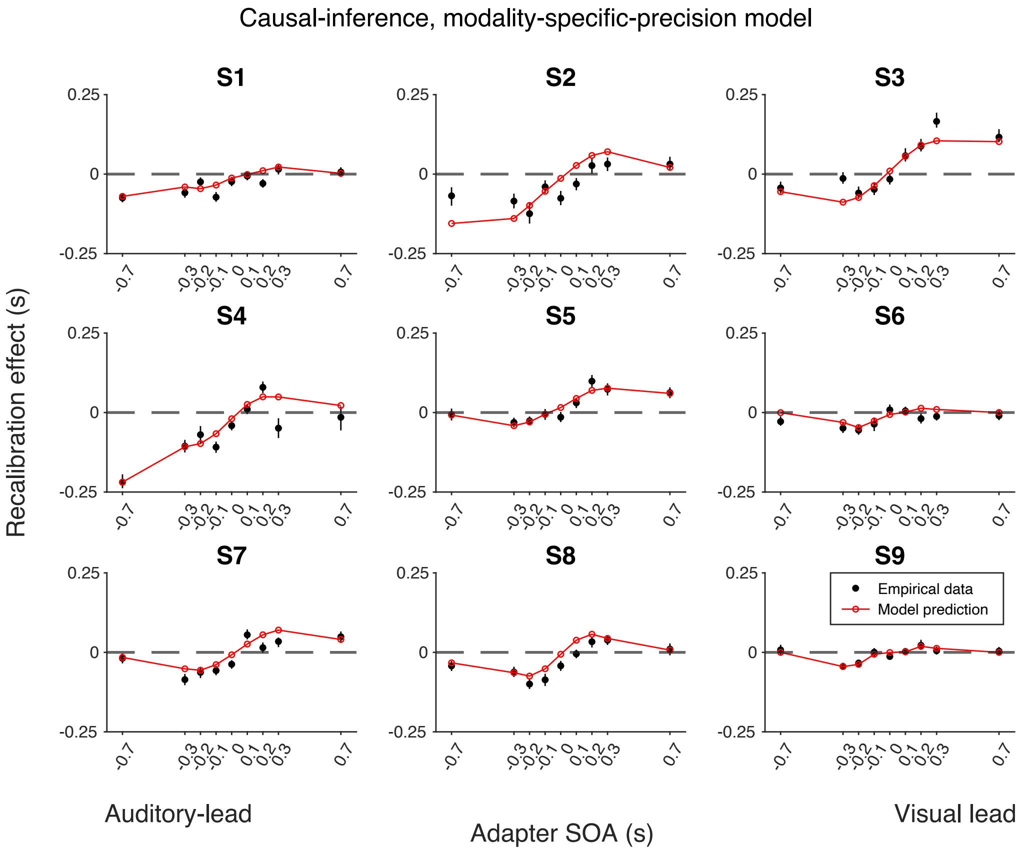

Figure S4: Empirical data for the recalibration effect for each participant and model predictions of the causal-inference recalibration model with modality-specific precision. Each panel represents a single participant. Black dots: empirical estimates from the best atheoretical model (the bias-shift model with exponential likelihood). Error bars: the 95% bootstrapped confidence intervals. Red lines: predictions of the recalibration model from the best-fitting parameter estimates for that participant.

#### 5.2 Asynchrony-contingent model

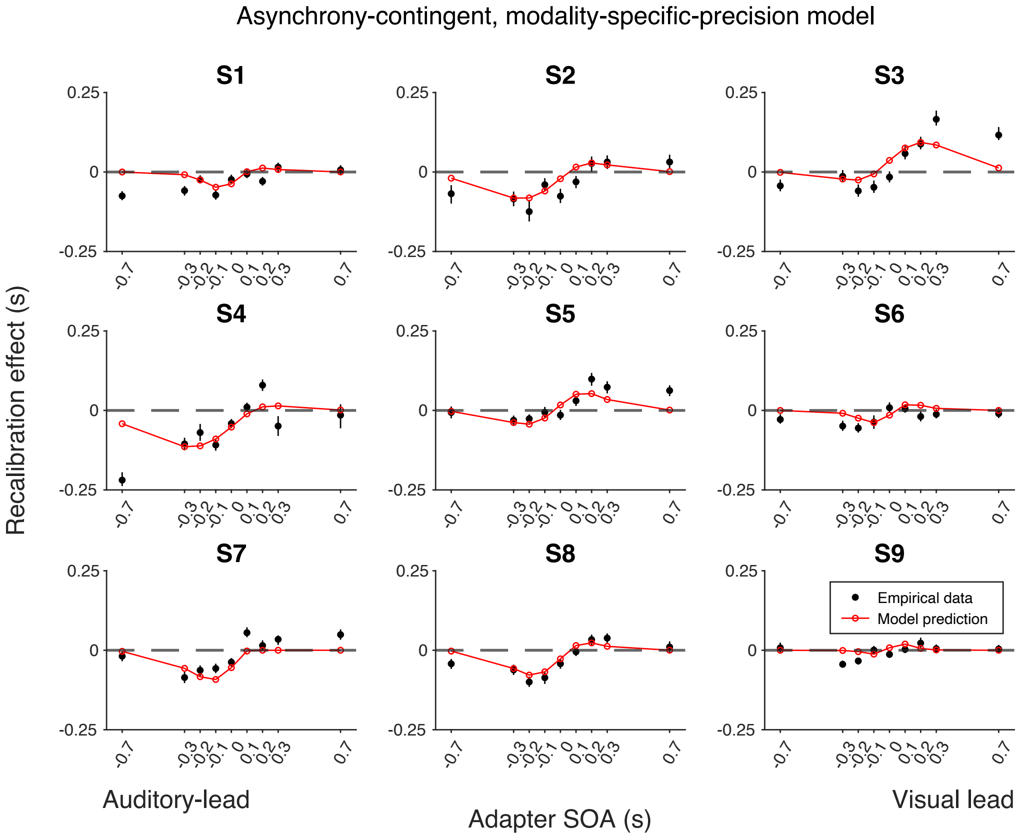

Figure S5: Empirical data of the recalibration effect for each participant and model predictions of the asynchrony-contingent recalibration model with modality-specific precision. Each panel represents a single participant. Black dots: empirical estimates from the best atheoretical model (the bias-shift model with exponential likelihood). Error bars: the 95% bootstrapped confidence intervals. Red lines: predictions of the recalibration model from the best-fitting parameter estimates for that participant.

#### 5.3 Asynchrony-correction model

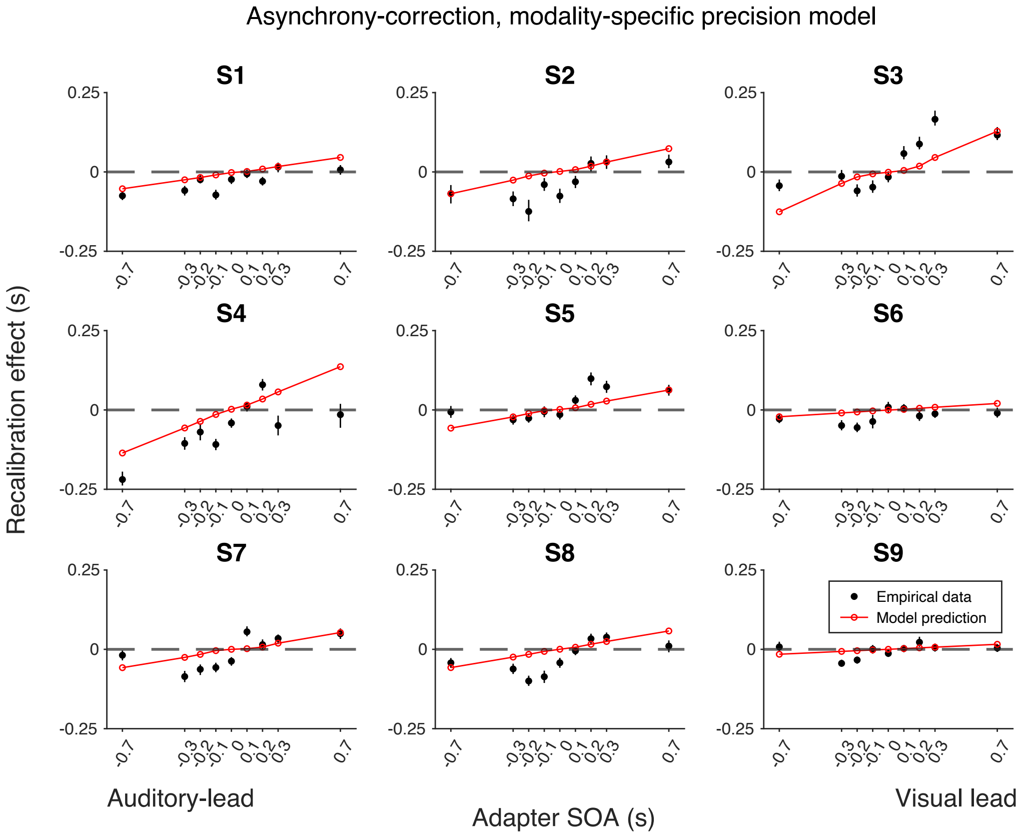

Figure S6: Empirical data of the recalibration effect for each participant and model predictions of the asynchrony-correction recalibration model with modality-specific precision. Each panel represents a single participant. Black dots: empirical estimates from the best atheoretical model (the bias-shift model with exponential likelihood). Error bars: the 95% bootstrapped confidence intervals. Red lines: predictions of the recalibration model from the best-fitting parameter estimates for that participant.

### 6 Scrutinizing the causal-inference model with modality-specific precision

#### 6.1 Individual parameter estimates

| | $\beta_{\text{pre}}$ | $\tau_A$ | $\tau_V$ | Criterion | $\lambda$ | $p_{common}$ | $\alpha$ | $\sigma_{C=1}$ | $\sigma_{C=2}$ |
|---|---|---|---|---|---|---|---|---|---|
| S1 | -66.78 | 49.76 | 67.89 | 91.74 | 0.005 | 0.671 | 0.003 | 69.51 | 214.24 |
| S2 | 45.86 | 90.41 | 144.48 | 185.86 | 0.047 | 0.498 | 0.007 | 70.66 | 390.54 |
| S3 | 10.94 | 94.48 | 86.64 | 190.74 | 0.008 | 0.504 | 0.007 | 40.05 | 260.65 |
| S4 | -4.42 | 80.78 | 113.14 | 105.57 | 0.054 | 0.221 | 0.014 | 17.99 | 267.67 |
| S5 | 30.15 | 89.35 | 66.38 | 109.35 | 0.049 | 0.571 | 0.004 | 69.53 | 263.09 |
| S6 | -9.79 | 41.69 | 62.88 | 85.54 | 0.029 | 0.345 | 0.006 | 69.71 | 528.98 |
| S7 | -31.37 | 68.79 | 60.78 | 59.69 | 0.051 | 0.864 | 0.002 | 18.57 | 218.73 |
| S8 | 0.16 | 66.37 | 76.78 | 63.25 | 0.017 | 0.378 | 0.008 | 26.48 | 247.67 |
| S9 | 14.03 | 33.22 | 41.23 | 47.12 | 0.014 | 0.632 | 0.007 | 73.70 | 575.44 |
| Mean | -1.25 | 68.32 | 80.02 | 104.32 | 0.031 | 0.520 | 0.006 | 50.69 | 329.67 |
| SEM | 11.10 | 7.51 | 10.41 | 17.32 | 0.007 | 0.064 | 0.001 | 8.16 | 45.53 |
| Lower bound | -200.00 | 10.00 | 10.00 | 1.00 | 0.000 | 0.000 | 0.000 | 1.00 | 100.00 |
| Upper bound | 200.00 | 200.00 | 200.00 | 350.00 | 0.060 | 1.000 | 0.020 | 300.00 | 1000.00 |

Table S1: Parameter estimates of individual participants. All parameters except $\lambda, p_{\text{common}}$ and $\alpha$ are in milliseconds.

#### 6.2 Model predictions of temporal-order judgments for individual participants

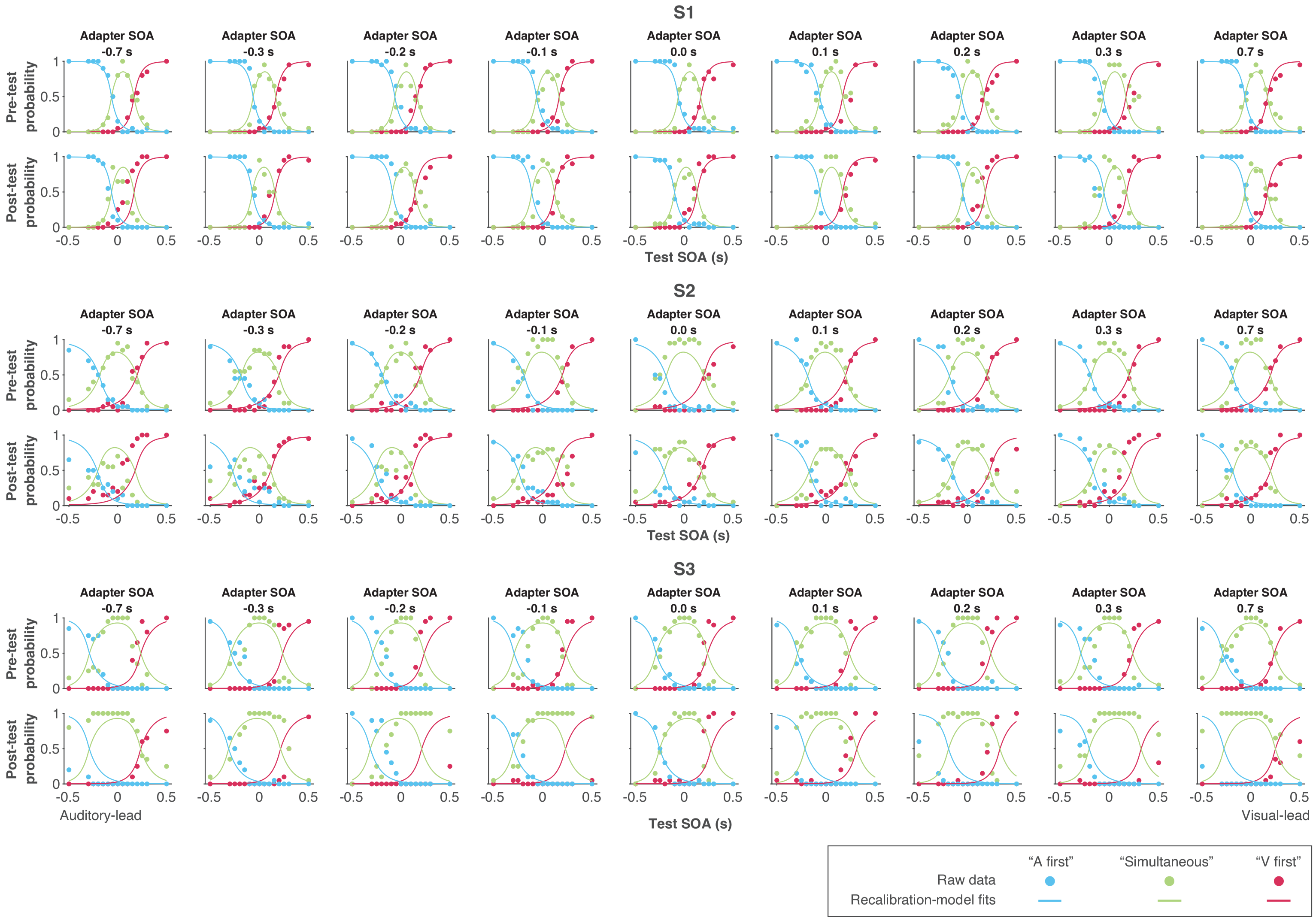

Figure S7: Data and predictions of the causal-inference, modality-specific-precision model for the temporal-order-judgment responses for each participant.

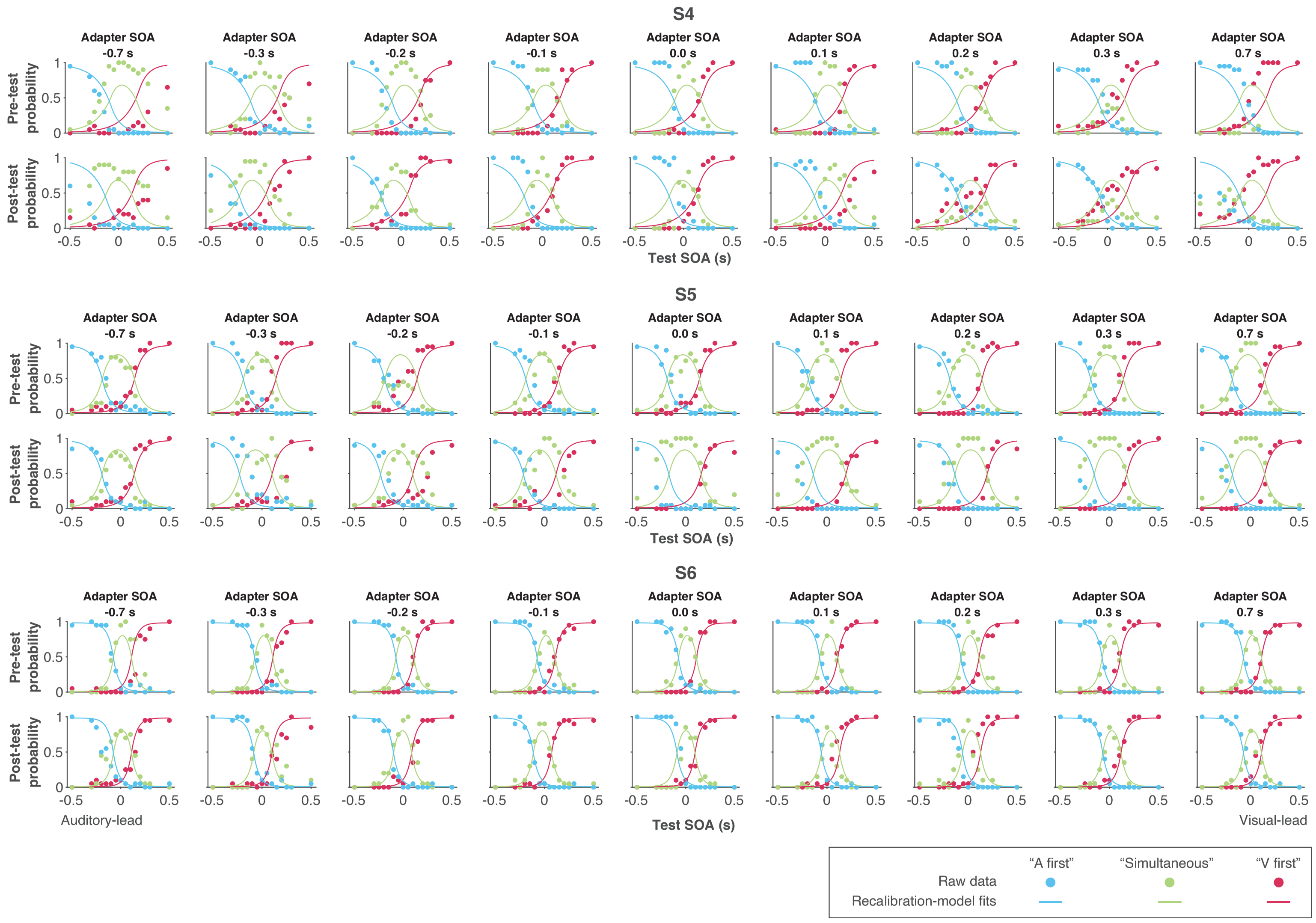

Figure S7: Data and predictions of the causal-inference, modality-specific-precision model for the temporal-order-judgment responses for each participant (continued).

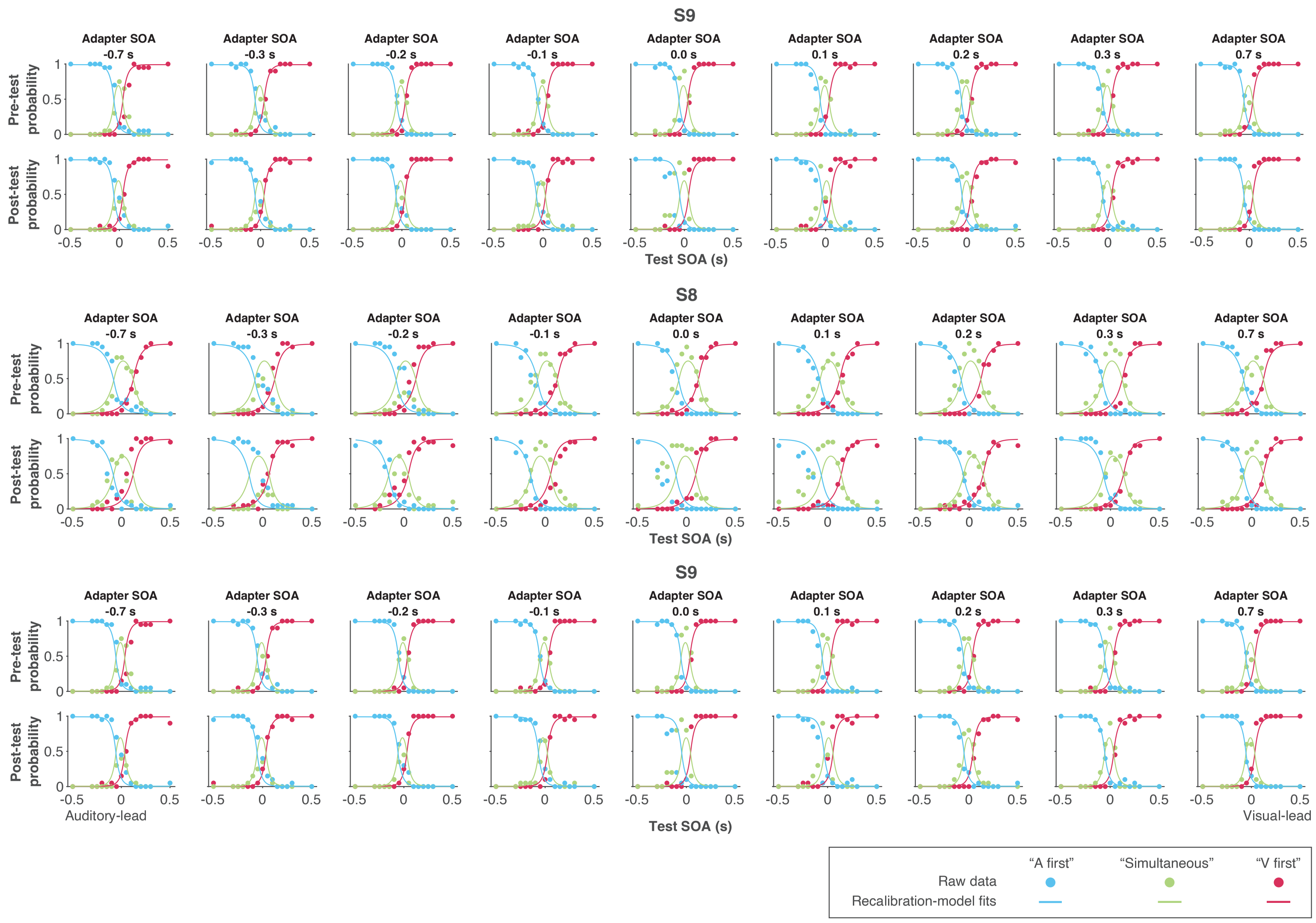

Figure S7: Data and predictions of the causal-inference, modality-specific-precision model for the temporal-order-judgment responses for each participant (continued).

### 7 Simulation of the causal-inference model

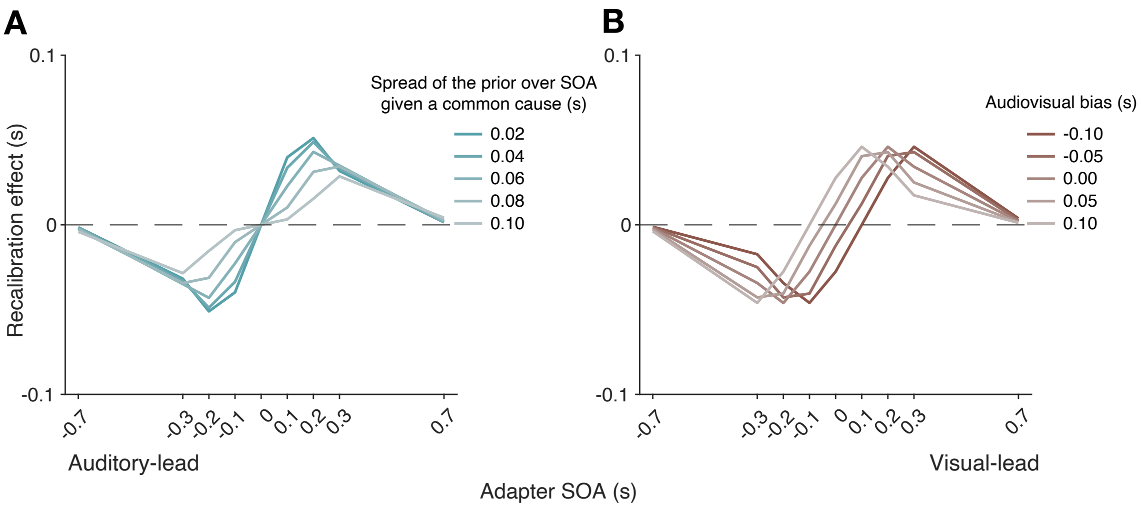

Figure S8: (A) The effect of the spread of the prior over SOA given a common cause on recalibration. A smaller spread increases the recalibration magnitude, but only for a small SOA range where the probability of a common cause is higher. (B) The effect of the audiovisual temporal bias on recalibration. Audiovisual temporal bias affects where recalibration starts in the direction of the other modality, but its effects are reduced for large adapter SOAs. Overall, temporal bias shifts the recalibration function laterally, with minimal impact on the asymmetry of recalibration effect.

### 8 Simulation of a population-code recalibration model

We simulated the population-code model of audiovisual temporal recalibration (Roach et al., 2011; Yarrow et al., 2015) to examine its prediction of the recalibration effect (i.e., the shift of the PSS) for different adapter SOAs. This model proposes that a population of neurons is tuned to different SOAs, and recalibration occurs through a selective gain reduction around the adapter SOA (Figure S9A, left panel). We assume a maximum-likelihood decoder that is unaware of the adaptation, resulting in a bias: estimates are shifted away from the adapted SOA (Figure S9A, middle panel; Jazayeri and Movshon, 2006; Seriès et al., 2009). The bias (i.e., the difference between estimated and physical SOA) is nonuniform as a function of the test SOA (Figure S9A, right panel).

When the adapter SOA is close to or exceeds the range of preferred SOAs across the neural population, it elicits less adaptation across neurons (Figure S9B, left panel) and results in a smaller change in population response (Figure S9B, middle panel). Consequently, the bias is significantly reduced as the adapter SOA increases (Figure S9B, right panel).

We simulated the bias in estimation as a function of stimulus SOA for this neural population when adapted to a range of different adapter SOAs (Figure S9C). These adapter SOAs produce a consistent non-uniform bias across stimulus SOAs. The overall magnitude of the bias is largest for small adapter SOAs.

Finally, we determined the recalibration effect by calculating the PSS after adaptation for each adapter SOA, assuming an unbiased PSS before adaptation (Figure S9D). That is, for each adapter SOA we determined the stimulus SOA that, after adaptation, resulted in an SOA estimate of 0 ms, that is, the stimulus SOA plus the bias is zero. The results show that the population-code model can predict nonlinearity in recalibration because the strength of adaptation decreases as the adapter SOA approaches the bounds of the range of preferred SOAs across the neural population. However, it cannot account for the asymmetry of recalibration observed in the empirical data, as neurons representing relative timing, a supramodal attribute, have similar selectivity in this model regardless of which modality leads.

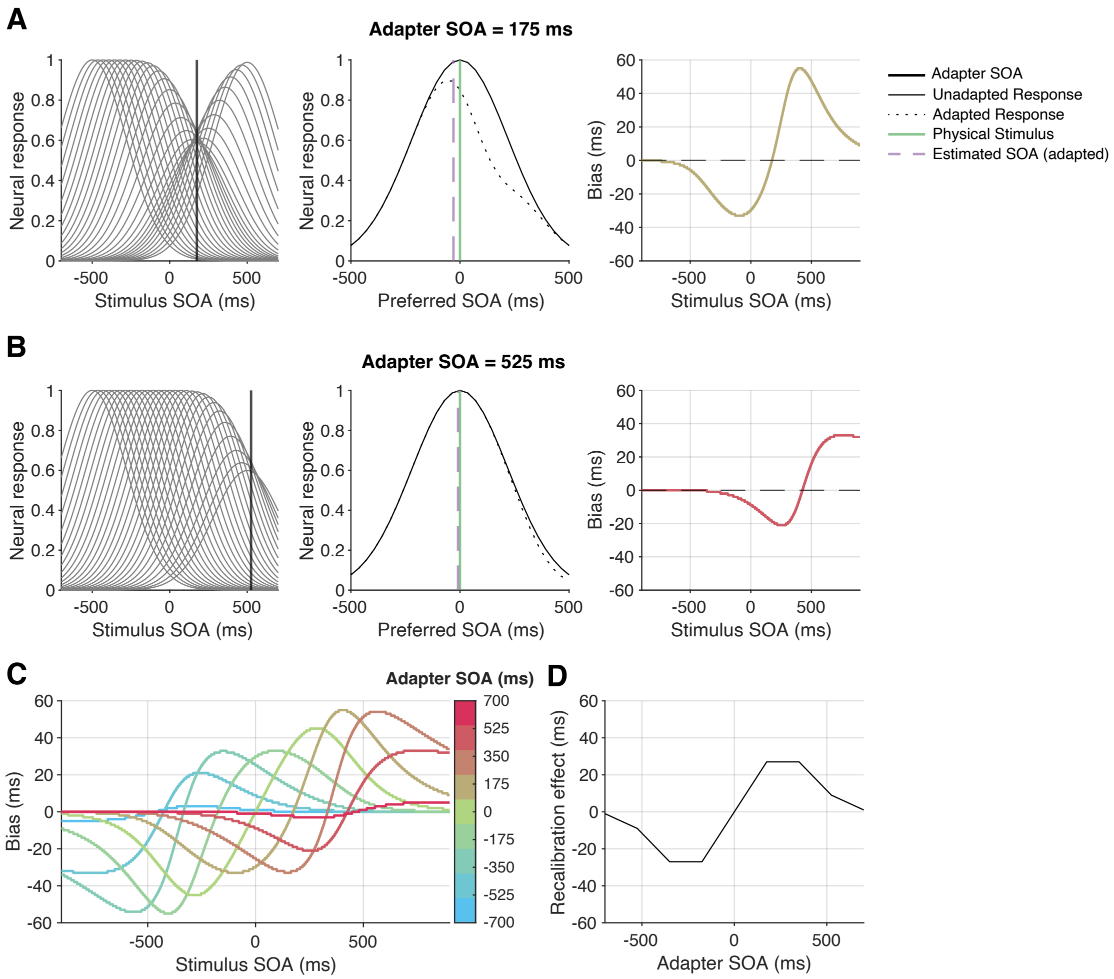

Figure S9: Simulation of a population-code model with varying adapter SOA. (A) Left panel: Tuning curves of a neural population adapted to a 175 ms SOA, with 29 neurons having preferred SOAs that range from -500 ms to 500 ms. Middle panel: neural responses to a stimulus with 0 ms SOA before (solid) and after (dashed) adaptation to a 175 ms SOA. Green: physical 0 ms SOA. Purple: estimated SOA based on an adaptation-unaware maximum-likelihood readout of the population response after adaptation. Right panel: Bias (i.e., estimated SOA minus physical SOA) across different stimulus SOAs after adaptation to a 175 ms SOA. (B) The same as in (A) after adaptation to a 525 ms SOA. (C) Bias as a function of stimulus SOA for several adapter SOAs. (D) Recalibration effect (i.e., the PSS: the stimulus SOA leading to an estimate of a 0 ms SOA) assuming an unbiased PSS before adaptation.

### 9 Exclusion of an outlier participant

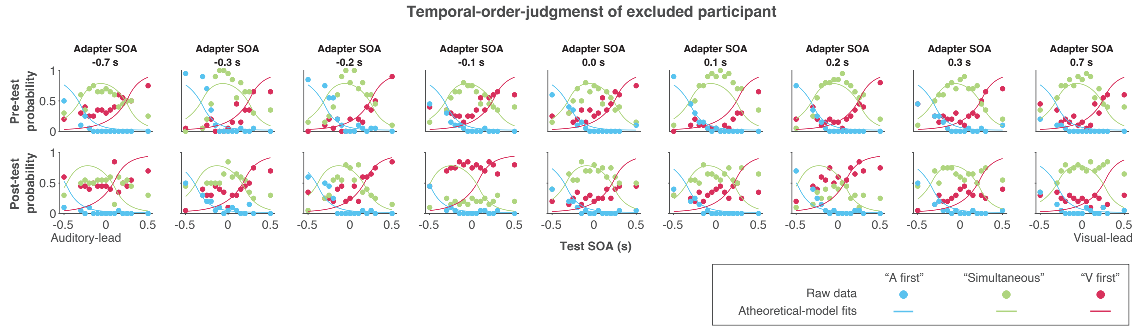

Figure S10: Temporal-order judgments from the outlier participant. We excluded this participant's data because the probabilities of the three responses were similar across test SOAs. These responses were too noisy to constrain the psychometric function.

#### 10 Performance in the oddball-detection task

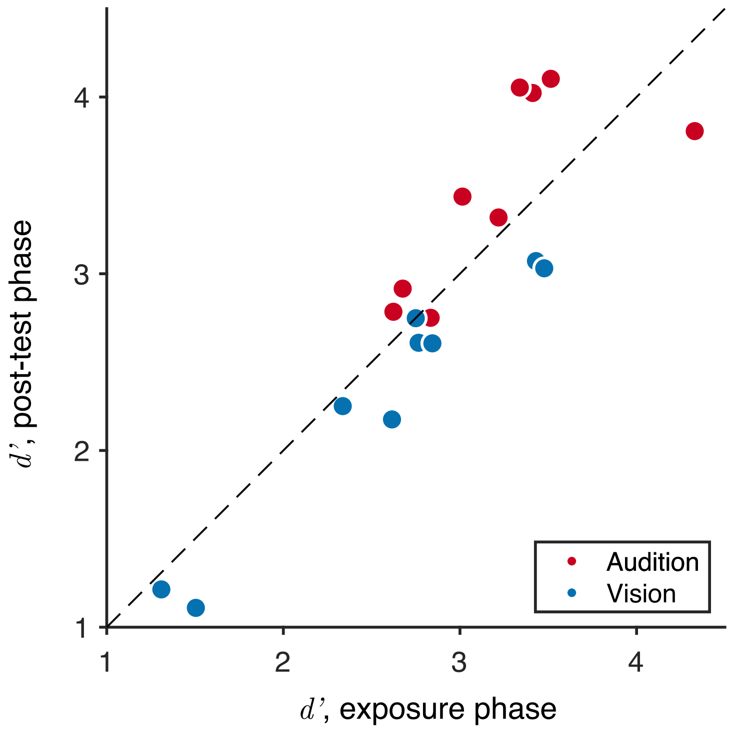

Figure S11: Oddball-detection performance in the exposure and post-test phases. Each dot represents a participant. In each phase, we combined responses across sessions and computed hit rates and false-alarm rates for auditory and visual oddballs. For each modality, hit and false-alarm rates were the probability of reporting an oddball when an oddball was presented and when it was absent, respectively. The trials in which both visual and auditory oddballs were presented were excluded when calculating hit and false-alarm rates, because participants did not indicate which modality the oddball was. The resulting $d'$ values for each participant confirm that they engaged in the task and attended to both modalities during the exposure phase. In general, participants showed a higher $d'$ for auditory oddballs compared to visual ones, despite our attempt to equate performance levels. Furthermore, the $d'$ remained consistent from the exposure to post-test phases, indicating similar performance in the top-up trials to performance during the exposure phase.

### 11 Model recovery of recalibration models

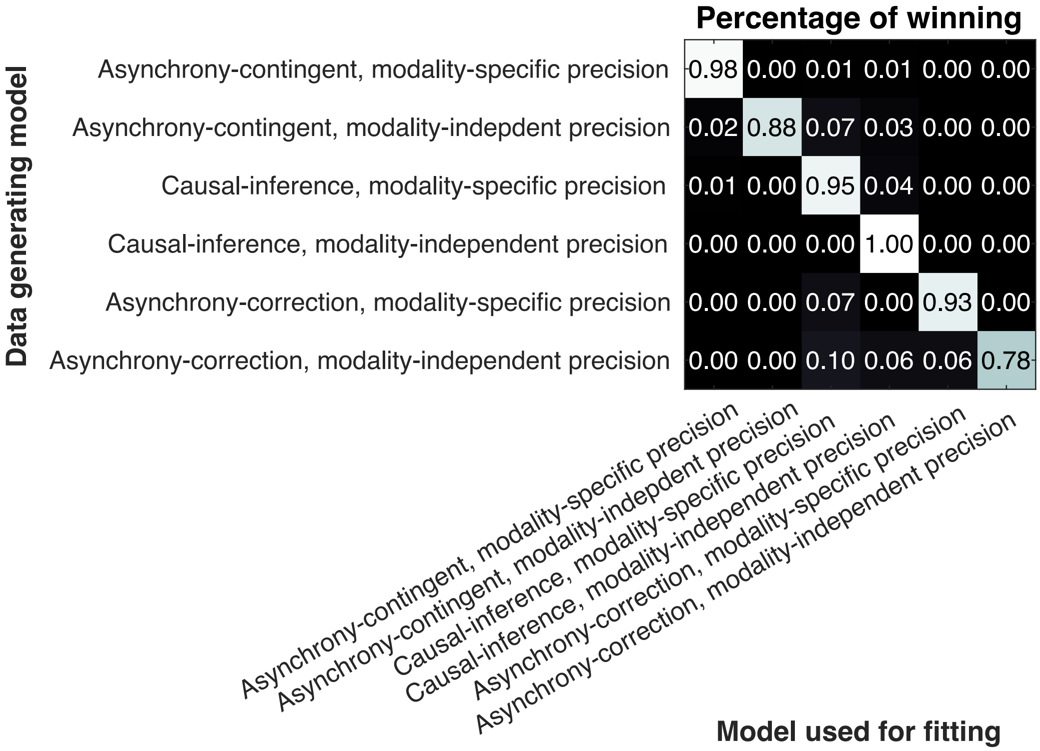

Figure S12: Confusion matrix resulting from the model-recovery simulations. Each cell indicates the proportion of the 100 simulations for each model generating the data that were best fit by each of the six models (i.e., the values in each row sum to one).

We conducted a model-recovery analysis for the six models described in the main text. In the main text, each model was fit to the data of each participant multiple times and the best-fitting parameter set for that model was selected. This yielded one parameter set for each participant and model. For each model, we randomly selected 100 parameter sets. Each parameter was randomly drawn from either a Gaussian (for unbounded parameters) or log-Gaussian (for parameters restricted to be positive) with mean and SD equal to that of the best-fit parameter sets across participants for that model. We then simulated datasets for each model based on each of the 100 parameter sets. All six models were fit to each simulated dataset.

We used the same model-comparison procedure as in the main text. All asynchrony-contingent and causal-inference models have satisfactory recovery results, suggesting that these models are identifiable. In addition, the data generated by the asynchrony-correction model were sometimes best fit by causal-inference models but not by the asynchrony-contingent models. This is reasonable since the asynchrony-correction model is similar to the causal-inference model with the value of $p_{\text{common}}$ set to one.

### 12 Model recovery of variants of the causal-inference recalibration models

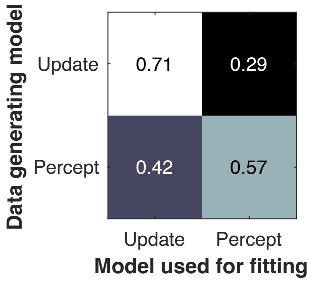

Figure S13: Confusion matrix resulting from the model-recovery simulations. Update model: the alternative model assuming that causal inference modulates recalibration at the bias-update phase. Percept model: the model described in the main text that assumes that causal inference modulates recalibration at the perceptual stage.

We compared the causal-inference recalibration model in the main text to a variant in which causal inference modulates recalibration in the update phase. Specifically, the learning rate is modulated by the posterior probability of a common cause, that is, the probability that the audiovisual signals arose from the same source. The bias update rule (Eq. 2) becomes:

$$\Delta_{\beta,i+1} = \Delta_{\beta,i} - P(C = 1|m_i)\alpha m_i. \tag{S7}$$

We performed a model-recovery analysis comparing these two models. 120 datasets were simulated for each model using the best-fitting parameters from the fit to the raw data. and we fit both models to each simulated dataset. The confusion matrix summarizes the probability that the generating model is the best-fitting model. Clearly, the results indicate that these two models are not distinguishable given the amount of data we collected in the main experiment.

### 13 Parameter recovery of the causal-inference, modality-specific-precision model

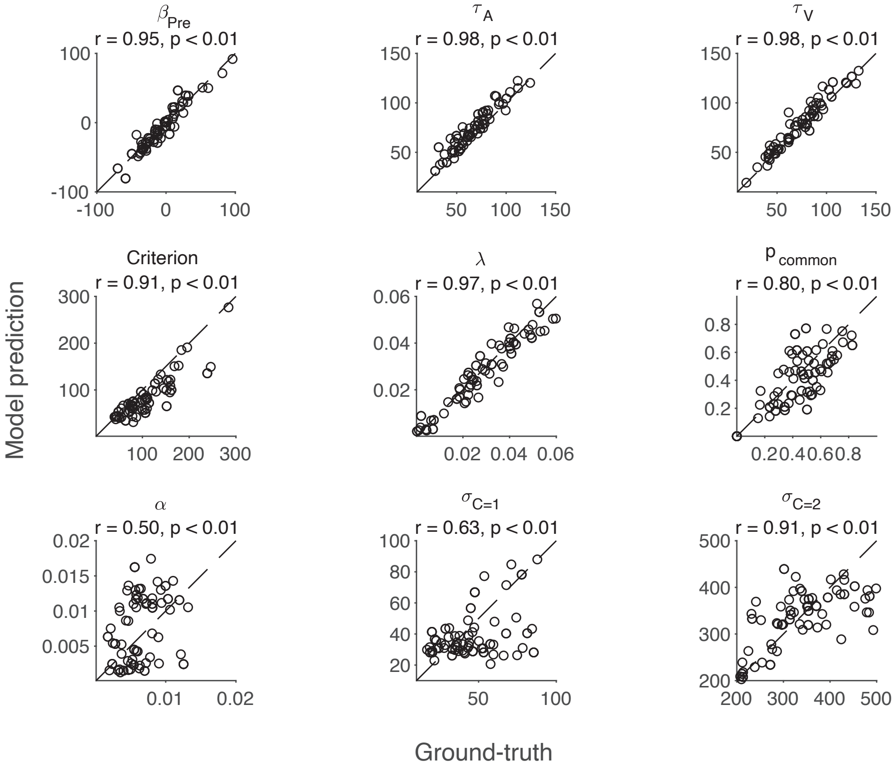

Figure S14: Predicted value vs. simulated value of each parameter. Each dot represents one out of 100 simulated datasets. The identity line marks perfect recovery performance.

We conducted a parameter recovery analysis of the winning model by simulating 100 datasets. Parameter values were sampled using the group mean and standard deviation across participants of the best-fitting estimates from model fits of the data. Each dataset was fit with the same causal-inference model with modality-specific precision. Key parameters, including audiovisual bias $\beta_{\text{pre}}$, amount of auditory latency noise $\tau_A$, amount of visual latency noise $\tau_V$, criterion, lapse rate $\lambda$ showed satisfactory recovery performance. The less accurate recovery of $p_{common}$ is likely due to a tradeoff with the learning rate $\alpha$, as both parameters influence the update of the audiovisual bias. The less accurate recovery of the widths of the SOA for a common cause ($\sigma_{C=1}$) and separate causes ($\sigma_{C=2}$) is acceptable, as their function can still be fulfilled within the respective range.
